# Reprogramming a Protein Ligase for Genetic Code Expansion

**DOI:** 10.64898/2026.07.07.736966

**Authors:** Giovanni Gallo, Alina Sieber, Michael Hellwig, Maximilian J. L. J. Fürst, Jürgen Lassak

## Abstract

The ribosome’s DNA-encoded production of defined polymer sequences is naturally limited to 22 amino acids. Although the translation machinery has the latent capacity to polymerize backbone-modified substrates, including β-amino acids, this potential is constrained by the intrinsic α-selectivity of native aminoacyl-tRNA synthetases. Here, we address this limitation by “reverse engineering” the *Escherichia coli* protein ligase EpmA. Naturally activating (*R*)-β-lysine, EpmA evolved to discard its tRNA-binding domain in favor of protein recognition. By grafting the anticodon-binding domain of the canonical lysyl-tRNA synthetase, LysRS, onto EpmA, we created the chimeric enzyme chEpmA. To our knowledge, this represents the first successful reprogramming of a protein ligase into a functional aminoacyl-tRNA synthetase. We demonstrate that chEpmA serves as a versatile dual-specificity platform: it efficiently charges tRNAs with the non-canonical backbone (*R*)-β-lysine, and a single substitution unlocks the scaffold for α-substrates, thereby enabling a broad spectrum of post-translational modifications previously inaccessible to genetic code expansion. This repertoire ranges from acylated lysines such as *N*_*ε*_*-*succinyl-(*S*)-α lysine (Ksucc) and bulky modifications such as biocytin to advanced glycation end products (AGEs) including *N*_*ε*_*-*carboxymethyl-(*S*)-α lysine (CML). Our work establishes a structural blueprint for mobilizing non-canonical substrates, paving the way for the biosynthesis of protease-resistant peptidomimetics and next-generation therapeutics.

---

**N**atural translation is limited to the polymerization of the 22 (*S*)-α-amino acids.^1^ While this conservation ensures high-fidelity protein synthesis, it restricts the chemical diversity of the resulting proteome. Expanding the genetic code to include backbone-modified substrates, specifically β-amino acids, is of particular interest for the development of peptidomimetics.^2-4^ Unlike canonical peptides, polymers containing β-amino acids, often referred to as foldamers, adopt unique secondary structures and exhibit intrinsic resistance to enzymatic degradation.^2,4^ Since endogenous proteases are evolved to recognize and cleave α-peptide bonds, the incorporation of β-backbones renders these molecules stable in biological environments, making them attractive candidates for next-generation peptide therapeutics with extended half-lives.^2,4^ However, the *in vivo* realization of this goal is especially hindered by the exquisite stereochemical selectivity of the native aminoacyl-tRNA synthetases (aaRSs) and the translation machinery.^5-7^ Recent studies have challenged the assumption that the ribosome cannot process β-backbones.^8-12^ On the one hand, *in vitro* translation experiments have demonstrated that the ribosome possesses a latent capacity for the co-translational polymerization of β-amino acids.^9,13,14^ On the other hand, powerful selection platforms such as tRNA display have enabled the evolution of pyrrolysyl-tRNA synthetase (PylRS) variants that charge tRNAs with β-amino acids.^11^ Despite these advances, the broad application of this technology *in vivo* is especially limited to aaRSs capable of efficiently charging tRNAs with β-amino acids. Canonical aaRSs are evolutionarily optimized for the α-backbone, creating a barrier to engineering β-specificity.^5,6^ To address this limitation, we focused on the bacterial protein ligase EpmA (YjeA, PoxA, GenX), a paralog of class II lysyl-tRNA synthetases (LysRS) (Scheme 1, left panel).^15,16^ In its physiological context, EpmA activates (*R*)-β-lysine with ATP to post-translationally modify translation elongation factor P (EF-P) (Scheme 1 right panel),^17-19^ a modification required to alleviate ribosomal stalling at proline-containing motifs.^20-22^ Evolutionarily, EpmA has retained the catalytic domain for amino acid activation, specifically adapting to the β-backbone^18^, but has lost the anticodon-binding domain required for tRNA recognition.^16,23^ Consequently, EpmA activates β-amino acids but transfers them to a protein substrate rather than tRNA.^24^ Here, we present a strategy to repurpose this specialized ligase into the two chimeric aminoacyl-tRNA synthetases, chEpmA-α and chEpmA-β, that enable tRNA charging with a wide variety of β- and α-amino acids, respectively, to either produce proteins altered at their backbone or co-translationally incorporate thus-far inaccessible post-translational modifications. Consequently, our work establishes chEpmA as a versatile enzymatic blueprint for the mobilization of non-canonical substrates for genetic code expansion.

**Scheme 1:**
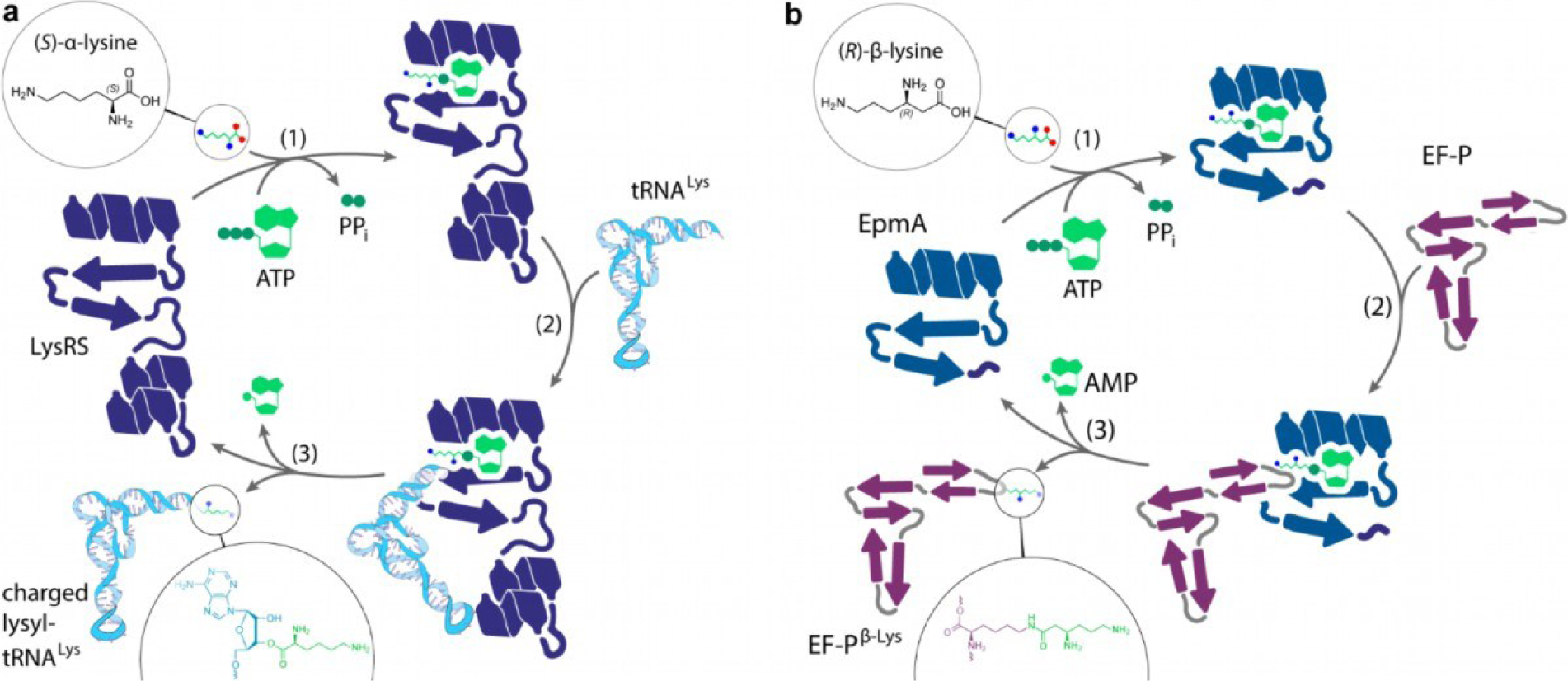
Mechanistic analogy between lysyl-tRNA charging by LysRS and *R-*β-lysylation of EF-P by EpmA. Left panel: LysRS catalyzes canonical aminoacylation by charging tRNA^Lys^ with (*S*)-α-lysine in an ATP-dependent reaction, yielding lysyl-tRNA^Lys^ and AMP. Right panel: the EF-P-modifying enzyme EpmA catalyzes post-translational β-lysylation of elongation factor P (EF-P) by transferring (*R*)-β-lysine to a specific acceptor site on EF-P, thereby consuming ATP and releasing AMP. Despite acting on distinct acceptors (tRNA versus protein), both enzymes proceed through the same three conceptual steps: **(1)** donor substrate activation via formation of an aminoacyl-adenylate intermediate, **(2)** ligation/transfer of the activated donor to the acceptor (the 3′ end of tRNA^Lys^ or EF-P), and **(3)** release of the charged acceptor (lysyl-tRNA^Lys^ or EF-P–β-Lys). Colored symbols denote the amino acid substrates (α-lysine *vs*. β-lysine) and ATP/AMP turnover, highlighting the shared chemical logic underlying amino acid transfer in translation and EF-P modification.

To transform the protein ligase EpmA into a functional aminoacyl-tRNA synthetase, we devised a strategy to create chimeras between the anticodon-binding domain (ABD) of the housekeeping LysRS LysS and EpmA. The critical challenge for these designs would be to correctly position the LysRS-derived relative to the EpmA catalytic core, and enable productive active site accommodation without compromising β activity. Before addressing this architectural problem, we first defined the catalytic framework that would guide both chimera design and subsequent functional validation. A central determinant of EpmA substrate specificity is the active-site residue A298, which acts as a backbone-selectivity gate. ^18,23^ In the native enzyme, A298 supports activation of (*R*)-β-lysine, whereas the A298G substitution reverses substrate preference towards (*S*)-α-lysine.^18,23^ We therefore considered two complementary chimeric scaffolds from the outset: the wild-type A298 version, hereafter referred to as chEpmA-β, designed to retain (*R*)-β-lysine specificity, and the corresponding A298G variant, chEpmA-α, designed to enable (*S*)-α-lysine-dependent tRNA charging. The latter variant was particularly important for functional validation *in vivo*, because direct testing of a β-lysine-specific chimera would be expected to charge the endogenous tRNA pool with β-lysine, causing proteome-wide misincorporation at canonical lysine codons and thereby imposing severe toxicity on the host (Fig. S14).

On this basis, we resorted to *in silico*-guided protein engineering to obtain a functional chimera that retained sufficient LysRS character to unlock tRNA activity while retaining EpmA-like chemistry. We first performed an exhaustive split-point (sp) scan along a carefully crafted mixed structure-and-sequence alignment of EpmA and LysRS. We systematically varied the crossover between the LysRS and the EpmA catalytic domain along the first 100 amino acids of the EpmA N-terminus that is equivalent to the domain linker in LysRS (Figure S1). To score each candidate chimera, we used Boltz-2 ^25^ to predict structures in six ligand configurations: apo, (*R*)-β-Lys-AMP, (*S*)-α-Lys-AMP, (*R*)-α-Lys-AMP, tRNA^Lys^ alone, and tRNA^Lys^ plus (*S*)-α-Lys-AMP co-bound – in both the A298 and A298G active-site, yielding approximately 13,700 structures in total (Figure 1, Figure S3). Models were then ranked using a composite score that integrated five criteria: structural integrity, ABD orientation, tRNA accommodation, catalytic-fold retention, and active-site competence. This analysis delineated a clear viability window between split-points sp15 and sp49, framed by N-terminal misfolding below sp10 and progressive erosion of β-specificity beyond sp50 (Figure 1). From this window, we selected sp43 as our lead design for chEpmA, because it combined a high composite score with good retention of the A298/A298G specificity switch. A thorough description of the computational workflow is provided as Supplementary Text. Having thus identified structurally plausible chimeras and established an α-lysine-specific version suitable for cellular testing, we next asked whether chEpmA-α could functionally operate as a genuine lysyl-tRNA synthetase *in vivo*. Specifically, we assessed whether chEpmA-α could functionally replace *E. coli* LysRS to sustain canonical protein synthesis under physiological conditions, by eliminating the two paralogs LysS and LysU.^26-28^ While the constitutively expressed LysRS LysS is responsible for housekeeping aminoacylation, the alternative LysU is described as heat-inducible.^26,29^ Nevertheless, *lysU* is expressed robustly under standard growth conditions and functionally complements the absence of LysS.^30^

**Figure 1:**
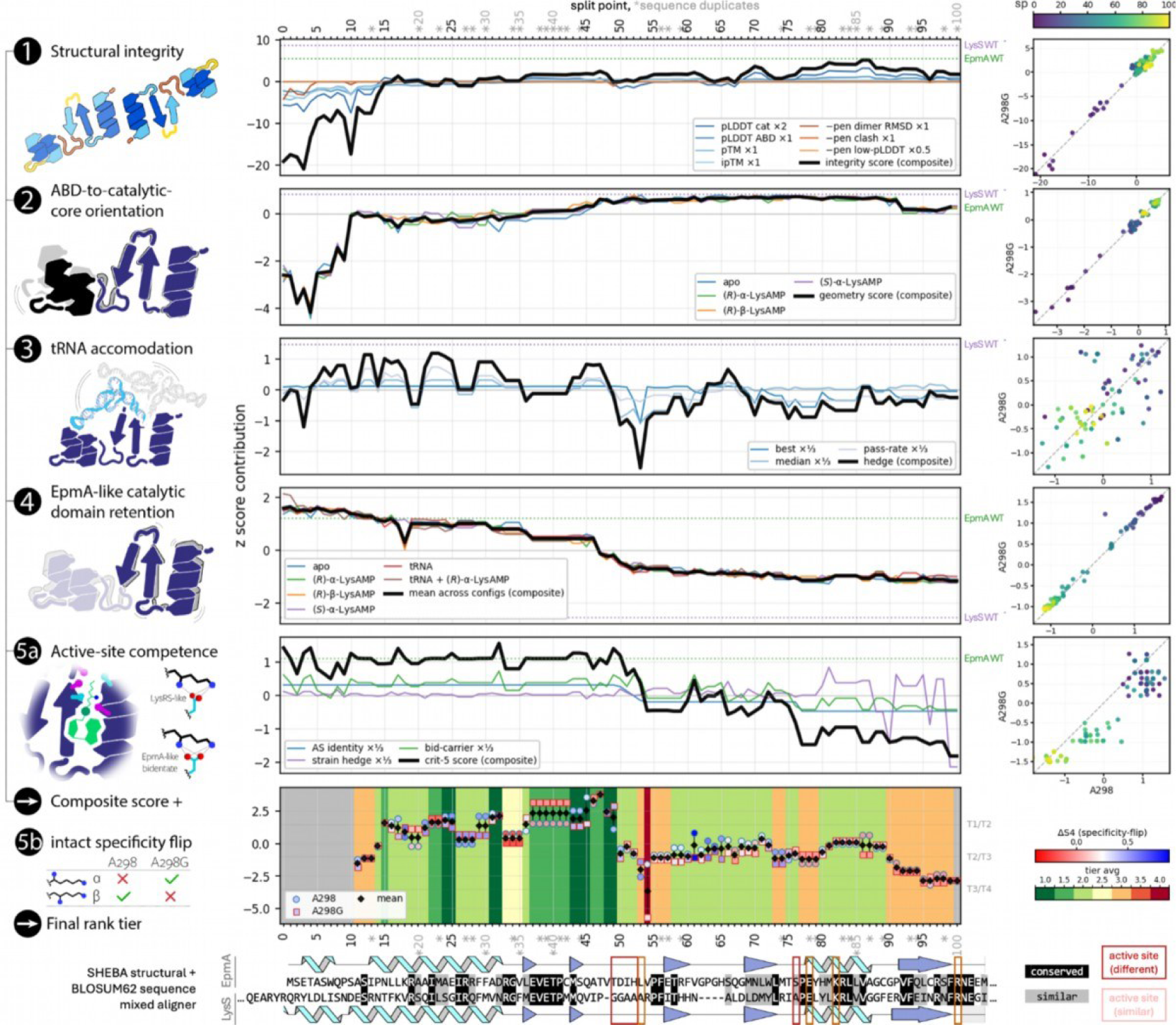
*In silico* screen of chimeras between LysRS anticodon-binding domain (ABD) and EpmA. To assess chimera candidates, we devised a computational structural modeling and analysis pipeline based on evaluating five structural modeling criteria **(left column)**. Different chimeras are characterized by a shifted split point (sp) along the N-terminus of EpmA, equivalent to a domain-bridging helix acting as a linker in LysRS **(bottom of the Figure)**. The chimeric sequences were obtained by moving the sp from position 0 (EpmA fully retained) to 100 (where the first 100 amino acids of EpmA are replaced by the equivalent residues in LysRS); where amino acid stretches with directly adjacent conserved residues cause neighboring sps to bear identical sequences (indicated with *). All unique chimeras were modeled with Boltz-2 and scored across the five metrics, yielding a composite score that integrated each criterion’s z-score **(central Figure column)**. The composite was then combined with an assessment of how well specificity for α vs β-Lys is retained in two active-site variants to create a final ranking across seven performance tiers. The criterion 1-5 plots show the data for the A298 variant, while the scatter plots (right column) show the limited divergence of the outcome for the A298G variant. See Supplementary Text for details and the exact definition of sub-criteria and legend shorthands.

We validated the functional integrity of our engineered scaffold using three parallel genetic strategies in an *E. coli* Δ*lysU* background (JW4090)^30^ (Figures 2a and S13). Starting with a strain lacking the auxiliary lysyl-tRNA synthetase ensures that cellular growth depends solely on the constitutive housekeeping function of LysS. First, we employed a CRISPR interference system^31^ to transcriptionally silence the remaining and now essential genomic *lysS* gene. While cells harboring an empty vector failed to propagate under knockdown conditions, expression of chEpmA-α fully rescued bacterial growth. Second, we demonstrated that the essential chromosomal *lysS* gene could even be deleted when chEpmA-α was provided *in trans* on a plasmid. At the same time, we made an interesting observation: during these plasmid-based complementation studies, high-level overexpression of EpmA_A298G, which encompasses only the catalytic core but lacks the grafted ABD, was also sufficient to rescue the lethal Δ*lysS* Δ*lysU* phenotype, albeit with reduced efficiency compared to the chimera. This finding points to a residual, intrinsic affinity of the EpmA wild-type enzyme for the tRNA acceptor stem, in turn implying that the “backbone barrier” might be less absolute than previously thought, thereby providing a latent evolutionary gateway for the stochastic entry of β-amino acids into the genetic code. Third and finally, to verify long-term genetic stability and prove that our chimera functions efficiently even without overexpression, we integrated the gene encoding chEpmA-α directly into the bacterial chromosome(Fig. S13e). In this background, we could successfully generate a Δ*lysS* Δ*lysU* double-deletion strain that relies exclusively on the genomically integrated chEpmA-α for survival. This strain exhibited robust growth characteristics, nearly comparable to those of the wild type, providing definitive proof that our rationally designed chimera functions as a housekeeping aminoacyl-tRNA synthetase. To substantiate our genetic findings with biochemical evidence and to determine the catalytic proficiency of the engineered synthetase, we performed *in vitro* aminoacylation assays. We purified *E. coli* LysS, wild-type EpmA, and the engineered chEpmA-α variant. A direct comparison of the steady-state kinetics highlights the functional impact of our rational design. We specifically determined the kinetic parameters for the tRNA substrate to quantify the efficiency of the grafted domain. Under our assay conditions, the native LysS exhibited high affinity for tRNA^Lys^, with a *K*_M_ of 2.1 ± 0.3 µM (Figure 2b)^32^. For our engineered chimera chEpmA-α we derived a *K*m for tRNA of 11.6 ± 2.8 µM. While this indicates a roughly 5-fold reduction in affinity compared to the natural housekeeping enzyme, it is crucial to place these values in a physiological context. The intracellular concentration of specific tRNA isoacceptors in *E. coli* typically ranges from 0.3 to 30 µM.^33^ Consequently, the Km of chEpmA lies well within the relevant physiological window, ensuring that the enzyme can efficiently charge tRNA *in vivo* despite the engineered interface. In terms of catalytic turnover, chEpmA-α showed a *k*_cat_ of 0.28 ± 0.03 min^−1^, compared to the isolated LysS of (1.8 ± 0.12 min^−1^) (Figure 2b). This confirms that the fusion of the heterologous ABD successfully re-establishes robust functional communication between the EpmA active site and the tRNA acceptor stem. By contrast, wild-type EpmA displayed only residual, albeit detectable tRNA charging activity under these conditions which was, however, too low to derive kinetic parameters. This invites us to speculate on the evolutionary forces that shaped the EpmA lineage. One plausible hypothesis is that the loss of the tRNA-binding domain was driven by negative selection that raised the dissociation constant (*K*_D_) above physiological tRNA concentrations. On the one hand, this may serve as a strict fail-safe mechanism to ensure the total exclusion of non-canonical backbones. On the other hand, it is tempting to hypothesize that the barrier is not absolute but rather permits a low level of sub-stoichiometric incorporation. Given that the ribosomal machinery is to some extent compatible with backbone-modified substrates,^12^ and that nature actively employs radical SAM enzymes to install β-amino acids post-translationally in proteins, it is conceivable that a co-translational route might also be evolutionarily preserved. In this scenario, the “backbone barrier” would function not as a wall, but as a tunable filter. Our data demonstrate that the EpmA catalytic core retains a fundamental geometric compatibility with the tRNA acceptor stem; whether this latent affinity is merely an evolutionary vestige or a functional feature allowing for stochastic proteomic diversification remains an open question.

**Figure 2:**
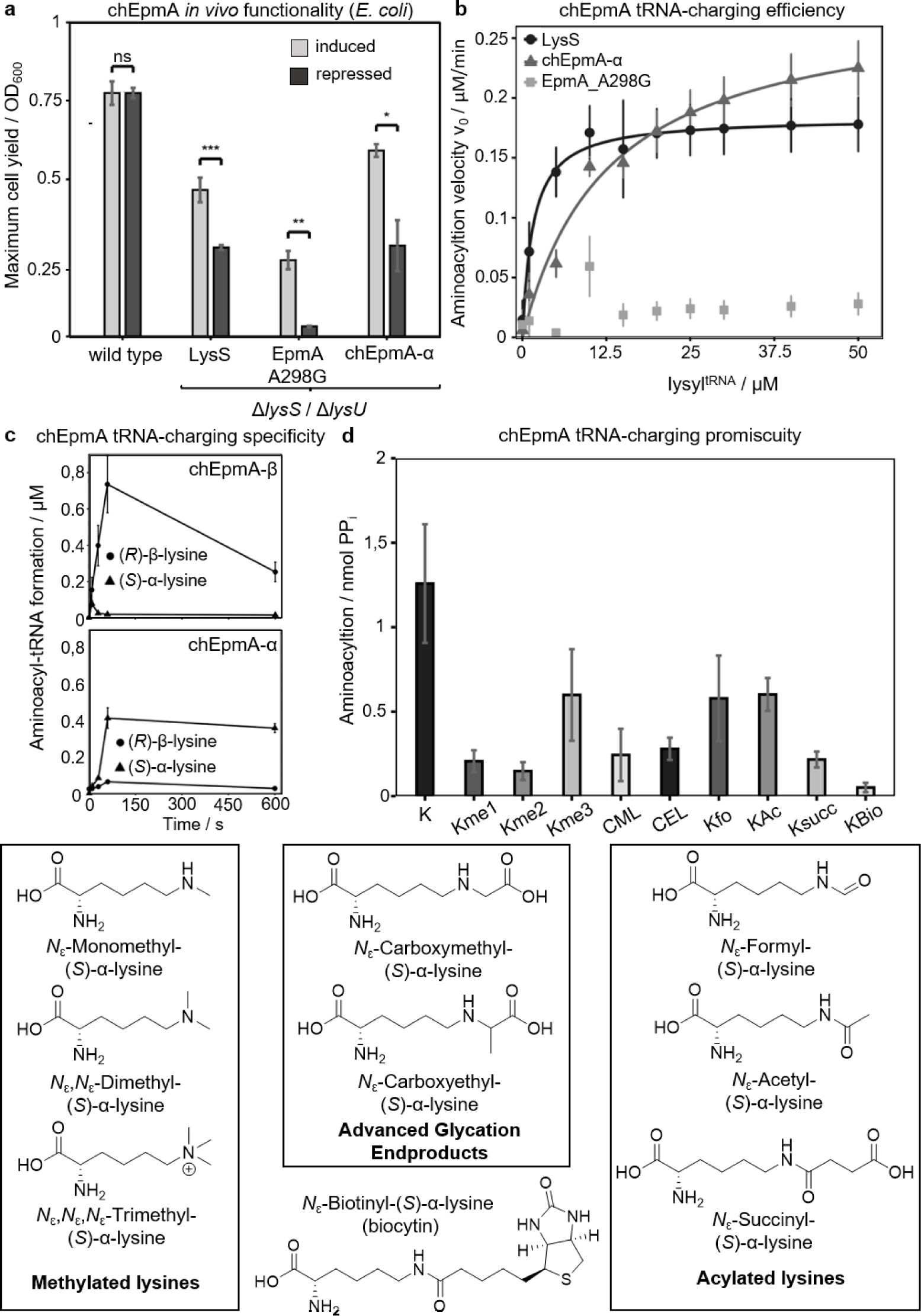
*In vivo* and *in vitro* validation of the engineered chEpmA synthetase and its α/β specificity switch. **a)** *In vivo* functional complementation: maximum cell yield (OD_600_) of *E. coli* Δ*lysS* Δ*lysU* carrying plasmid-borne LysRS, EpmA_A298G, or chEpmA-α, recorded after 16 h of growth under arabinose-induced (0.2% w/v) versus glucose-repressed (0.2% w/v) conditions; wild type denotes the parental strain carrying the empty vector. Statistical significance was assessed by unpaired two-tailed Student’s *t*-test (ns, not significant; *p < 0.05; **p < 0.01; ***p < 0.001). The full set of complementation strategies (CRISPRi, plasmid complementation and genomic integration) is shown in Figure S13. **b)** *In vitro* steady-state kinetics for tRNA^Lys^ charging with L-α-lysine at saturating concentration (1 mM), using 1 µM enzyme; initial rates were determined from the amount of aminoacylated product formed after 10 min. chEpmA-α and EpmA_A298G compared with *E. coli* LysRS as the canonical benchmark. **c)** Time-course aminoacylation of chEpmA-β (upper panel) and chEpmA-α (lower panel) with 1 µM enzyme and 1 mM (*R*)-β-lysine or 1 mM (*S*)-α-lysine. **c)** Substrate spectrum of chEpmA-α for tRNA^Lys^-charging with a variety of physiologically relevant *N*ε-modified L-lysine derivatives. Reactions contained 1 µM enzyme and 1 mM of the indicated amino acid substrate, and were incubated for 16 hours at 37 °C; tRNA aminoacylation activity (Amount of PPi / nmol) was quantified as described in the Experimental Section. Abbreviations for substrates: K, (*S*)-α-lysine; Kme1, *N*_ε_-monomethyl-(S)-α-lysine; Kme2, *N*ε,*N*ε-dimethyl-(*S*)-α-lysine; Kme3, *N*ε,*N*ε,*N*ε-trimethyl-(*S*)-α-lysine; CML, *N*ε-carboxymethyl-(*S*)-α-lysine; CEL, *N*ε-carboxyethyl-(*S*)-α-lysine; Kfo, *N*_ε_-formyl-(*S*)-α-lysine; Kac, *N*_ε_-acetyl-(*S*)-α-lysine; Ksucc, *N*_ε_-succinyl-(*S*)-α-lysine; KBio, Biocytin, *N*_ε_-biotinyl-(*S*)-α-lysine. All values are baseline-corrected by subtracting the corresponding tRNA^Lys^-only control (no amino acid substrate; null line). Error bars indicate variability between replicates.

Having established that the engineered chEpmA-α variant functions as a proficient canonical synthetase, we sought to determine if this chimeric scaffold could be “switched back” to its original specificity, effectively creating a dedicated module for the β-world. To define this orthogonality, we performed time-course aminoacylation assays, comparing the charging profiles of the glycine substitution A298G (chEpmA-α) and the variant with the restored wild-type alanine A298 at the active site (chEpmA-β) against the native LysS (Figure 2c). The results reveal the expected functional dichotomy. The chEpmA-α variant behaved indistinguishably from the housekeeping LysRS: both enzymes showed rapid, linear charging of tRNA^lys^ with (*S*)-α-lysine while remaining completely inert towards (*R*)-β-lysine. In stark contrast, reverting glycine to the wild-type alanine (chEpmA-β) resulted in a complete inversion of specificity. This variant exhibited robust aminoacylation exclusively with (*R*)-β-lysine, with no detectable activity towards the α-substrate. This confirms that the single residue at position 298 acts as a binary switch,^18^ allowing us to toggle the enzyme’s selectivity between the two backbone worlds without compromising the restored tRNA affinity. Crucially, this clean “toggle” highlights the privileged nature of the EpmA scaffold. While we demonstrate here that the EpmA architecture can be effortlessly converted into a canonical synthetase, the reciprocal engineering of LysRS towards β-lysine specificity appears significantly more restricted: An equivalent single mutation in *Bacillus cereus* LysRS, which has a strong similarity to the active site of EpmA (also see SI text section Figure S6), does not confer robust activity towards the β-substrate.^34^ This functional asymmetry indicates that β-backbone recognition is governed not by a single residue but by the overall architecture of the EpmA catalytic core. Consequently, our chEpmA scaffold represents a unique platform: it is the only known enzymatic architecture that can be readily tuned to charge tRNA with both α- and β-amino acids. With our two well-defined variants in hand, we sought to explore the substrate scope beyond the binary distinction of (*S*)-α-versus (*R*)-β-lysine. Previous investigations by our laboratory and others have collectively highlighted a remarkable substrate plasticity of the EpmA catalytic core.^15,23,35^ These studies demonstrated that the enzyme accepts a surprisingly broad range of lysine derivatives. Most notably, McDonnell *et al*. revealed that the native enzyme tolerates “extreme” modifications, including β-lysine analogs with extended alkyl and aminoalkyl chains (up to C10-decyl)^35^. Furthermore, the catalytic core exhibits remarkable stereochemical and chemical versatility, accepting hydroxylated side chains and even the non-natural (*S*)-configuration at the β-position.^18,23^ This proves that the distal region of the active site can accommodate significant hydrophobic and polar bulk, as well as distinct stereochemistries, without significantly compromising catalysis. Based on these robust data, we propose a refined evolutionary hypothesis: the ancestral “gatekeeping” mechanism acted strictly as a molecular sieve defined by only two geometric parameters, the minimal chain length and the precise positioning of the β-amine. Since the discrimination against the canonical proteome relied almost exclusively on these backbone features, there was likely minimal evolutionary pressure to constrain the remainder of the substrate-binding pocket.

We applied these insights into the active site architecture to test the potential of our engineered scaffold. By using the chEpmA-α variant, we effectively bypassed the backbone filter, allowing us to probe the active site’s tolerance to side-chain modifications on a canonical α-amino acid scaffold *in vitro* (Figure 2d). We challenged the enzyme with a panel of physiologically relevant substrates that mimic important post-translational modifications. The enzyme exhibited robust tRNA-charging activity with *N*_ε_-acetyl-(*S*)-α-lysine (Kac) and Ksucc, confirming its ability to accommodate both charged and uncharged modifications. Most notably, chEpmA-α also accepted the AGEs CML and *N*_ε_-carboxyethyllysine (CEL), and even the sterically demanding biocytin can serve as a weak off-target. These findings confirm that the substrate pocket is exceptionally spacious and largely agnostic to the nature of the ε-modification. Consequently, our work provides a blueprint for a modular enzymatic platform where backbone specificity (α vs. β) and side-chain tolerance are effectively decoupled. By simply toggling the steric gate while exploiting the pocket’s inherent promiscuity, this scaffold offers a rational strategy to access the entire coordinate system of lysine modifications, ranging from post-translational marks to pathological AGEs, using a single, structurally unified enzyme family. Our study establishes the chEpmA scaffold as the functional blueprint for the enzymatic incorporation of beta-amino acids into peptides and proteins. To advance this proof-of-concept into a fully orthogonal “cell factory,” we propose a streamlined engineering roadmap that builds upon already validated physiological adaptations. The first step, decoupling the system from host survival, has already been achieved. Since (*R*)-β-lysine is inherently essential for the native EpmA, we previously replaced the endogenous elongation factor machinery with the orthogonal arginine-rhamnosylation pathway from pseudomonads^16^. This established “system swap” maintains cell viability while releasing β-lysine from its housekeeping duties, effectively rendering it an orthogonal metabolite. The next step addresses translational orthogonality. To target free codons parallel to the genetic code, the synthetase requires an anticodon-binding domain. Analogous to the chimeric strategy recently described by Ding *et al*., we propose fusing the N-terminal domain of the orthogonal synthetase PylRS to the chEpmA core.^36^ When paired with a chimeric tRNA, this creates a dedicated translation channel without cross-reactivity. Implementation of these steps will yield a fully autonomous *E. coli* host capable of the total biosynthesis of proteins containing beta-amino acids. While the engineering of uptake systems is undeniably a valuable contribution towards industrial scalability,^37^ endogenous biosynthesis represents the method of choice. Our approach leverages this distinct advantage: by generating substrates directly within the cytoplasm, using aminomutases to convert abundant precursors, we bypass external transport entirely. This ensures a continuous, self-sufficient, and stereochemically precise supply of building blocks. Importantly, this metabolic module is highly versatile: replacing the *E. coli* mutase EpmB with its clostridial homolog, KamA, switches production from (*R*)-to (*S*)-β-lysine,^38^ while recruiting specific acetyltransferases enables access to side-chain-modified variants such as β-acetyl-lysine.^39^ Ultimately, this unlocks two transformative applications. First, in the realm of therapeutics, the site-specific incorporation of cationic, protease-resistant beta-backbones offers a decisive solution to the stability limitations of host-defense peptides (HDPs), paving the way for a new class of durable antibiotics.^40,41^ Second, the remarkable plasticity of the chEpmA active site opens a new frontier in biomedical research. We demonstrated that the binding pocket is sufficiently promiscuous to accommodate physiologically relevant side chains, including AGEs such as CML and CEL.^42-44^ These modifications, which typically accumulate stochastically during aging, may soon be installed site-specifically. By enabling the precise introduction of individual AGEs, this capability provides a unique tool for novel bottom-up investigations into the structural and functional consequences of protein glycation, achieving a level of precision currently unattainable with standard methods.

## Supporting Information

The authors have cited additional references within the Supporting Information.^[30, 31]^

## Acknowledgements

We thank the Boehringer Ingelheim Foundation for funding through the Exploration Grants Program (project “Genetic code expansion by β-amino acids”). We are also grateful to Thomas Carell for providing succinyl-lysine. The computational work used the Dutch national e-infrastructure with the support of the SURF Cooperative, using grant no. EINF-4326. We also thank the Center for Information Technology of the University of Groningen for providing access to the Hábrók high-performance computing cluster. We thank Jingzhi Stritzel for fruitful discussions.

## References

1 de la Torre, D. & Chin, J. W. Reprogramming the genetic code. Nat Rev Genet 22, 169–184 (2021). 10.1038/s41576-020-00307-7

2 Scott, T. A. et al. Widespread microbial utilization of ribosomal β-amino acid-containing peptides and proteins. Chem 8, 2659–2677 (2022). 10.1016/j.chempr.2022.09.017

3 Dong, N. et al. Short symmetric-end antimicrobial peptides centered on β-turn amino acids unit improve selectivity and stability. Front Microbiol 9, 2832 (2018). 10.3389/fmicb.2018.02832

4 Gopalan, R. D., Del Borgo, M. P., Mechler, A. I., Perlmutter, P. & Aguilar, M. I. Geometrically precise building blocks: the self-assembly of beta-peptides. Chem Biol 22, 1417–1423 (2015). 10.1016/j.chembiol.2015.10.005

5 Woese, C. R., Olsen, G. J., Ibba, M. & Söll, D. Aminoacyl-tRNA synthetases, the genetic code, and the evolutionary process. Microbiol Mol Biol Rev 64, 202–236 (2000). 10.1128/mmbr.64.1.202-236.2000

6 Rubio Gomez, M. A. & Ibba, M. Aminoacyl-tRNA synthetases. Rna 26, 910–936 (2020). 10.1261/rna.071720.119

7 Agirrezabala, X. & Frank, J. From DNA to proteins via the ribosome: structural insights into the workings of the translation machinery. Hum Genomics 4, 226–237 (2010). 10.1186/1479-7364-4-4-226

8 Melo Czekster, C., Robertson, W. E., Walker, A. S., Soll, D. & Schepartz, A. In vivo biosynthesis of a β-amino acid-containing protein. J Am Chem Soc 138, 5194–5197 (2016). 10.1021/jacs.6b01023

9 Katoh, T. & Suga, H. Ribosomal incorporation of consecutive β-amino acids. J. Am. Chem. Soc. 140, 12159–12167 (2018). 10.1021/jacs.8b07247

10 Fujino, T., Goto, Y., Suga, H. & Murakami, H. Ribosomal synthesis of peptides with multiple beta-amino acids. J Am Chem Soc 138, 1962–1969 (2016). 10.1021/jacs.5b12482

11 Dunkelmann, D. L. et al. Adding α,α-disubstituted and β-linked monomers to the genetic code of an organism. Nature 625, 603–610 (2024). 10.1038/s41586-023-06897-6

12 Cruz-Navarrete, F. A. et al. β-Amino Acids Reduce Ternary Complex Stability and Alter the Translation Elongation Mechanism. ACS Central Science 10, 1262–1275 (2024). 10.1021/acscentsci.4c00314

13 Lee, J. et al. Ribosomal incorporation of cyclic β-amino acids into peptides using in vitro translation. Chemical Communications 56, 5597–5600 (2020). 10.1039/D0CC02121K

14 Katoh, T., Tajima, K. & Suga, H. Consecutive elongation of D-amino acids in translation. Cell Chem Biol 24, 46–54 (2017). 10.1016/j.chembiol.2016.11.012

15 Ambrogelly, A., O’Donoghue, P., Söll, D. & Moses, S. A bacterial ortholog of class II lysyl-tRNA synthetase activates lysine. FEBS Lett 584, 3055–3060 (2010). 10.1016/j.febslet.2010.05.036

16 Lassak, J. et al. Arginine-rhamnosylation as new strategy to activate translation elongation factor P. Nat. Chem. Biol. 11, 266–270 (2015). 10.1038/nchembio.1751

17 Navarre, W. W. et al. PoxA, YjeK, and elongation factor P coordinately modulate virulence and drug resistance in Salmonella enterica. Mol Cell 39, 209–221 (2010). https://doi.org/S1097-2765(10)00462-4 [pii] 10.1016/j.molcel.2010.06.021

18 Roy, H. et al. The tRNA synthetase paralog PoxA modifies elongation factor-P with (R)-β-lysine. Nat Chem Biol 7, 667–669 (2011). 10.1038/nchembio.632

19 Yanagisawa, T., Sumida, T., Ishii, R., Takemoto, C. & Yokoyama, S. A paralog of lysyl-tRNA synthetase aminoacylates a conserved lysine residue in translation elongation factor P. Nat Struct Mol Biol 17, 1136–1143 (2010). https://doi.org/nsmb.1889 [pii] 10.1038/nsmb.1889

20 Ude, S. et al. Translation elongation factor EF-P alleviates ribosome stalling at polyproline stretches. Science 339, 82–85 (2013). 10.1126/science.1228985

21 Doerfel, L. K. et al. EF-P is essential for rapid synthesis of proteins containing consecutive proline residues. Science 339, 85–88 (2013). 10.1126/science.1229017

22 Lassak, J., Stritzel, J., Schlundt, A. & Brewer, T. Constant trouble with prolines—navigating a global translation dilemma. Nucleic Acids Research 54 (2026). 10.1093/nar/gkaf1420

23 Pfab, M. et al. Synthetic post-translational modifications of elongation factor P using the ligase EpmA. FEBS J 288, 663–677 (2021). 10.1111/febs.15346

24 Lassak, J., Sieber, A. & Hellwig, M. Exceptionally versatile take II: post-translational modifications of lysine and their impact on bacterial physiology. Biol Chem (2022). 10.1515/hsz-2021-0382

25 Passaro, S. et al. Boltz-2: Towards Accurate and Efficient Binding Affinity Prediction. bioRxiv (2025). 10.1101/2025.06.14.659707

26 Clark, R. L. & Neidhardt, F. C. Roles of the two lysyl-tRNA synthetases of Escherichia coli: analysis of nucleotide sequences and mutant behavior. Journal of Bacteriology 172, 3237–3243 (1990). 10.1128/jb.172.6.3237-3243.1990

27 Chen, X. et al. Multiple catalytic activities of Escherichia coli lysyl-tRNA synthetase (LysU) are dissected by site-directed mutagenesis. Febs j 280, 102–114 (2013). 10.1111/febs.12053

28 Onesti, S. et al. Structural studies of lysyl-tRNA synthetase: conformational changes induced by substrate binding. Biochem 39, 12853–12861 (2000). 10.1021/bi001487r

29 Emmerich, R. V. & Hirshfield, I. N. Mapping of the constitutive lysyl-tRNA synthetase gene of Escherichia coli K-12. J Bacteriol 169, 5311–5313 (1987). 10.1128/jb.169.11.5311-5313.1987

30 Baba, T. et al. Construction of Escherichia coli K-12 in-frame, single-gene knockout mutants: the Keio collection. Mol Syst Biol 2, 2006.0008 (2006). 10.1038/msb4100050

31 Qi, L. S. et al. Repurposing CRISPR as an RNA-guided platform for sequence-specific control of gene expression. Cell 152, 1173–1183 (2013). 10.1016/j.cell.2013.02.022

32 Brevet, A., Chen, J., Lévêque, F., Blanquet, S. & Plateau, P. Comparison of the enzymatic properties of the two Escherichia coli lysyl-tRNA synthetase species. J Biol Chem 270, 14439–14444 (1995). 10.1074/jbc.270.24.14439

33 Dong, H., Nilsson, L. & Kurland, C. G. Co-variation of tRNA abundance and codon usage in Escherichia coli at different growth rates. J Mol Biol 260, 649–663 (1996). 10.1006/jmbi.1996.0428

34 Gilreath, M. S. et al. β-Lysine discrimination by lysyl-tRNA synthetase. FEBS Lett 585, 3284–3288 (2011). https://doi.org/S0014-5793(11)00673-9 [pii] 10.1016/j.febslet.2011.09.008

35 McDonnell, C. M., Ghanim, M., Mike Southern, J., Kelly, V. P. & Connon, S. J. De-novo designed β-lysine derivatives can both augment and diminish the proliferation rates of E. coli through the action of elongation factor P. Bioorg Med Chem Lett 59, 128545 (2022). 10.1016/j.bmcl.2022.128545

36 Ding, W. et al. Chimeric design of pyrrolysyl-tRNA synthetase/tRNA pairs and canonical synthetase/tRNA pairs for genetic code expansion. Nat Commun 11, 3154 (2020). 10.1038/s41467-020-16898-y

37 Iype, T. et al. Hijacking a bacterial ABC transporter for genetic code expansion. Nature 647, 1045–1053 (2025). 10.1038/s41586-025-09576-w

38 Behshad, E. et al. Enantiomeric free radicals and enzymatic control of stereochemistry in a radical mechanism: the case of lysine 2,3-aminomutases. Biochem 45, 12639–12646 (2006). 10.1021/bi061328t

39 Müller, S. et al. Bacterial abl-like genes: production of the archaeal osmolyte N_ε_-acetyl-β-lysine by homologous overexpression of the yodP-kamA genes in Bacillus subtilis. Appl Microbiol Biotechnol 91, 689–697 (2011). 10.1007/s00253-011-3301-8

40 Ting, D. S. J., Beuerman, R. W., Dua, H. S., Lakshminarayanan, R. & Mohammed, I. Strategies in translating the therapeutic potentials of host defense peptides. Front Immunol 11 (2020). 10.3389/fimmu.2020.00983

41 Porter, E. A., Weisblum, B. & Gellman, S. H. Mimicry of Host-Defense Peptides by Unnatural Oligomers: Antimicrobial β-Peptides. Journal of the American Chemical Society 124, 7324–7330 (2002). 10.1021/ja0260871

42 Delgado-Andrade, C. Carboxymethyl-lysine: thirty years of investigation in the field of AGE formation. Food Funct 7, 46–57 (2016). 10.1039/c5fo00918a

43 Ulrich, P. & Cerami, A. Protein glycation, diabetes, and aging. Recent Prog. Horm. Res. 56, 1–21 (2001). 10.1210/rp.56.1.1

44 Ashraf, J. M. et al. Recent advances in detection of AGEs: Immunochemical, bioanalytical and biochemical approaches. IUBMB Life 67, 897–913 (2015). 10.1002/iub.1450

