## Supplementary Information for "Reprogramming a Protein Ligase for Genetic Code Expansion"

[1] Ludwig-Maximilians-Universität München
Fakultät für Biologie
Großhaderner Str. 2-4; D-82152 Planegg-Martinsried; Germany
[2] Technische Universität Dresden
Professur für Spezielle Lebensmittelchemie
Bergstraße 66; D-01069 Dresden; Germany
[3] University of Groningen
Molecular Enzymology
Feringa Building, Groningen, 9747AG, Netherland
[4] University of Naples Federico II
Department of Agricultural Sciences
Piazza Carlo di Borbone 1; 80055; Portici; Italy

#### S1. Engineering rationale

As a step toward enzymatically charging tRNA with β-amino acids and ultimately ribosomal synthesis of β-peptides, we set out to graft the anticodon-binding domain (ABD) of *E. coli* lysyl-tRNA synthetase (LysRS) onto the catalytic core of its β-lysine-activating homolog EpmA, which has evolutionarily lost the ABD. The central engineering choice was the position along the EpmA/LysRS alignment at which to insert the fusion: every viable design had to retain the entire ABD, but the exact crossover (or *split point*) would determine how the appended domain interfaces with the EpmA catalytic core. A poor junction might prevent the assembled protein from folding, while a split reaching too far into either the ABD or the EpmA catalytic domain would disrupt tRNA binding or β-lysine specificity, respectively. We approached this question computationally, predicting chimera dimer structures across a comprehensive split-point scan in two active-site contexts and six ligand configurations, and ranked the predicted models with a multi-criterion structural assessment to nominate candidates for experimental kinetic characterization.

#### S2. Defining split point 0 and the sweet-spot hypothesis

We first defined split point 0 (sp 0) as the alignment position marking the C-terminal boundary of the LysS ABD, the LysS region absent from EpmA, and therefore the minimal LysS segment that every chimera in our candidate pool had to carry (Figure S1). As the fusion site, sp 0 yields a marginal chimera: the full EpmA chain with the ABD grafted onto its N-terminus and no other LysS residues. This is the construct one would build under the simplest evolutionary assumption, that EpmA lost its ABD outright while leaving its N-terminus geometrically unaltered. We considered this assumption too simplistic: the EpmA N-terminus, freed of the constraint of supporting an ABD, has plausibly drifted into new conformations or acquired EpmA-specific roles. If so, sp 0 grafts the ABD onto an EpmA stretch that is no longer prepared to act as its linker, thereby perturbing inter-domain geometry and forfeiting any contribution the native LysS linker region might make to tRNA accommodation. Provided the EpmA N-terminus is itself dispensable for β-lysine activation, either functionally irrelevant or recruited to bind EpmA's evolutionarily new acceptor, EF-P, extending the LysS portion past sp 0 should replace the drifted EpmA N-terminus and restore a LysS-like linker. Assuming further LysS encroachment past this linker would begin to erode β-lysine specificity, our goal was to identify an expected "sweet spot" at intermediate split points.

#### S3. Hybrid structure–sequence EpmA–LysS alignment and chimera construction

Residue equivalence between homologs is ultimately defined by occupying the same position in 3D space. A pure sequence-based alignment of EpmA and LysS is inadequate for the present purpose: the EpmA N-terminus has weak homology to its LysS counterpart, allowing sequence aligners to open large gaps early in the chain and re-anchor only on the catalytic core, leaving sp 0 essentially indeterminate; further along, several indel-bearing internal stretches mislead aligners into pairing structurally non-equivalent residues, producing chimeras with spurious indels or non-equivalent crossover sites. We therefore built the EpmA–LysS alignment in two layers. As the primary source of truth, we aligned the EpmA and LysS catalytic-domain crystals with SHEBA, which anchored 248 structurally equivalent residue pairs (~72 % of the resulting 346-column alignment) at 1.17 Å Cα-RMSD. The remaining ~28 %, residues unresolved in one or both crystals, plus pairs that lie too far apart in 3D for SHEBA to call structurally equivalent, were filled with a BLOSUM62 pairwise sequence alignment carrying an extra terminal-gap penalty to favor flush end alignment. Most positions in the resulting map are high-confidence; the remaining low-confidence residues sit in regions that are simultaneously unresolved in crystals and poorly conserved in sequence, most notably the ~11 N-terminal residues of EpmA, which adopt a distinct conformation from their LysS equivalents and are an integral part of the corresponding catalytic-to-ABD linker in LysS.

The resulting alignment was used to programmatically generate every chimera in the split-point sweep: at split point n, the chimera consists of the full LysS ABD followed by LysS residues for alignment columns 0…n−1, and EpmA residues for columns n…end. The construction spans sp 0 (the full EpmA chain with the LysS ABD appended) to sp 345 (essentially full-length LysS with only the C-terminal-most EpmA residue retained). Indels in the alignment produce two side-effects: at an EpmA-deletion column, moving from sp n to sp n+1 adds one LysS residue while the EpmA portion is unchanged; at an EpmA-insertion column, the LysS portion is unchanged, but EpmA loses one residue, leaving that chimera one residue shorter than its alignment-position neighbors. Where the two enzymes share runs of identical residues, adjacent chimeras carry identical sequences and were collapsed to a single canonical split point downstream (Figure S2).

#### S4. Inventory of ground-truth structures

To validate the chimera scoring against known biology, we surveyed the public PDB for crystal structures of EpmA, LysS, and their class-II aaRS homologs. Our priority was the E. coli enzymes, the proteins on which we would build the chimeras, but in parallel, we collected homologous structures across the class-II aaRS fold to provide structural examples of Lys-AMP donor and tRNA acceptor binding modes. We combined a Foldseek search against the PDB with an RCSB GraphQL query spanning EC number, gene-name match, and ≥ 30% sequence identity to E. coli homologs, then filtered for false positives and consolidated the results into a master table of 607 candidate entries. E. coli EpmA and LysS are represented by multiple structures in apo and ligand-bound states, but two important complexes are absent from the public databases: no class-II representative has been crystallized with (R)-β-Lys-AMP, and no bacterial LysS–tRNA complex has been solved, leaving the human cytosolic LysS structure (PDB 9DPL) as the only tRNA-bound reference at this fold.

#### S5. A five-criterion structural scoring framework

For a chimera to function as intended, we reasoned it had to perform well along five complementary axes: (1) **dimer integrity**: does the homodimer hold together as a coherent fold; (2) **inter-domain orientation**: does the appended ABD sit in a pose capable of delivering a docked tRNA to the active site; (3) **tRNA accommodation**: can a $tRNA^{Lys}$ actually dock at the catalytic site in a geometrically productive orientation; (4) **catalytic-fold retention**: does the catalytic core remain structurally EpmA-like despite carrying LysS-derived residues at its upstream end; and (5) **active-site competence**: can the active site bind *(R)*-β-Lys-AMP in a chemically reasonable geometry and with an EpmA-like pocket character. Criteria 1–2 test whether the chimera is a viable protein at all; criteria 3–5 test whether, given a viable scaffold, it has the structural ingredients required for the desired activity on the intended substrate. The resulting computational pipeline is depicted in Figure S3. For each criterion, the corresponding panel of **main text Fig. 1** decomposes the score into its constituent submetrics so that the contribution of each submetric to the criterion score is visible along the split-point sweep; the definitions and rationale of those submetrics are spelled out below.

**Criterion 1: dimer integrity.** Criterion 1 combines Boltz-2's model-typical confidence metrics with structural sanity checks, averaged over the n = 10 ensemble per chimera. Four positive components contribute: the mean pLDDT over the catalytic-domain residues (pLDDT cat, weight 2, the catalytic core is the engineering substrate), the ABD-domain mean pLDDT (pLDDT ABD, weight 1), the chain-level pTM (weight 1), and the homodimer-interface ipTM (weight 1). Three penalties are subtracted: a chain-A-vs-chain-B Cα-RMSD penalty (pen dimer RMSD, weight 1) flagging asymmetric protomers, a steric-clash penalty (pen clash, weight 1), and a longest-low-pLDDT-catalytic-stretch penalty (pen low-pLDDT, weight 0.5), which catches localised disorder that the catalytic-domain mean pLDDT averages over. The seven components are z-scored relative to the cohort and summed to form the criterion-1 composite, which is also the integrity-gate score (see *Composite ranking*, S10).

**Criterion 2: inter-domain orientation.** Criterion 2 quantifies whether the LysS-derived ABD sits in a pose compatible with its expected resting orientation relative to the catalytic core, as established by the human cytosolic LysS chain A from the LysS–tRNA complex (PDB 9DPL). For each chimera and each of the four ligand-only configurations: apo, (*R*)-β-LysAMP, (*R*)-α-LysAMP, and (*S*)-α-LysAMP, the prediction is structurally aligned over its catalytic-domain Cα atoms to the reference, and the Cα-RMSD of the ABD residues is computed against the 9DPL ABD over a fixed residue-correspondence map. The per-config decomposition in the main text, Fig. 1, covers only these four ligand-only configurations because preliminary inspection showed that the tRNA-bearing predictions rarely converged on a coherent tRNA docking pose, leaving too few productively docked models per cell to support stable per-config statistics. The criterion-2 composite (the geometry_score) is the pooled mean of per-model ABD-RMSDs across the four ligand-only configurations and across the few tRNA-bound models that pass a per-model tRNA-dock coherence check; the latter admission shifts the per-cell score by ≤ 0.07 Å (within the wild-type noise floor). The four per-config Cα-RMSDs are individually z-scored across the cohort (sign-flipped so that lower RMSD yields higher z); decomposition by configuration makes ligand-induced ABD perturbations directly visible.

**Criterion 3: tRNA accommodation.** Criterion 3 evaluates whether a docked $tRNA^{Lys}$ is geometrically positioned for productive aminoacylation, applied to the two tRNA-containing configurations. For each Boltz model, the chimera–tRNA complex is structurally aligned to the human LysRS–tRNA reference (PDB 9DPL), and a per-model productivity score is computed as the negative sum of four geometric penalty terms: a sigmoid on the tRNA CCA-end-to-active-site distance above 9 Å, a sigmoid on the anticodon-loop-to-ABD-anchor distance above 6 Å, a linear penalty on the CCA-arm-to-anticodon-arm angle cosine below the wild-type-reference mean, and a quadratic penalty on the rigid-body elbow dihedral outside the wild-type-reference window. Models with productivity score > −10 are considered passing (the wild-type chimera-reference complex sits at ≈ 0). The n = 40 ensemble per cell is aggregated into three submetrics shown in *main text Fig. 1* as best (highest-scoring model), median (typical model), and pass-rate (fraction of models above the threshold). Each is z-scored against the cohort, and the criterion-3 composite is their unweighted mean, a *hedge* that balances best-case, typical-case, and robustness signals against the three failure modes the criterion guards against. The ×⅓ notation in main text Fig. 1 reflects this construction: each submetric is plotted at one-third of its z-magnitude so that the visual sum equals the composite hedge curve.

**Criterion 4: catalytic-fold retention.** Criterion 4 quantifies the EpmA-likeness of the chimera's catalytic-domain Cα geometry. The catalytic-domain Cα atoms are independently superposed onto the EpmA and LysS catalytic-domain crystal references, and the per-residue Cα-Cα distance to each parent is tabulated. The scalar $\delta = \Sigma_i (d_{i,LysS} - d_{i,EpmA})$, summed over catalytic-domain residues, is positive when the chimera is closer to EpmA than to LysS and negative when it has drifted toward LysS. Unlike criterion 2, criterion 4 is evaluated for all six ligand configurations: the catalytic-domain fold should remain EpmA-like in every ligand state. Each per-config δ is z-scored across the cohort (sign-flipped so that EpmA-like maps to higher z), and the criterion-4 composite is their unweighted mean across configurations.

**Criterion 5: active-site competence.** Criterion 5 was the most demanding to formalize: a complete structure-function model must not only define what a favorable ligand-binding mode is, but also stereoselectively discriminate productive from unproductive poses, despite EpmA being known to bind its non-cognate substrate (S)-α-Lys and despite cofolding models' tendency to pack similar ligands into known pockets in ways that may not reflect biological reality. The derivation of the criterion-5 rules from crystal-anchored geometry and substrate-kinetic evidence is the subject of S6; in the criterion-5 panel of the main text Fig. 1, the composite decomposes into three scored sub-scores, AS identity, strain hedge, and bid-carrier, together with a fourth, pose-in-pocket, reported as a sanity-check diagnostic.

#### S6. Defining criterion 5: linking active-site geometry to substrate kinetics

To establish the rules for criterion 5, we analyzed crystal structures of class-II aaRS active sites alongside the published kinetic evidence on substrate acceptance. The only intermediate-bound EpmA structure is in complex with an (S)-α-Lys-AMP analog, providing a non-cognate-substrate pose anchor; no *E. coli* LysS–Lys-AMP structure exists, but the 89 % identical *E. coli* LysU paralog has been crystallized with (S)-α-Lys-AMP, allowing us to infer the productive α mode. A second anchor is the EpmA A298G mutant, which weakens β activity and enhances α acceptance, demonstrating that a single sub-Ångström-scale change at position 298 can switch substrate preference. The EpmA and LysS active-site pockets are otherwise highly conserved, differing principally at position 298 (Gly in LysS), at Ser76/Ala239, and at a loop reorganization in which LysS residues 216–218 fold on top of the active site while the equivalent EpmA loop flips outward.

To benchmark Boltz-2 against this body of evidence, we co-folded all combinations of LysS, EpmA, and EpmA A298G with (*S*)-α-Lys and *(R)*-β-Lys substrates and systematically measured catalytic-anchor distances, per-residue pocket displacements vs. crystal, and the internal-coordinate strain of each ligand against PDB-pooled reference distributions, aggregated over the n = 10 ensemble. The analysis yielded a single robust separator: a paralog-architecture asymmetry at the catalytic glutamate (Glu116 in EpmA, Glu279 in LysS). In LysS the sidechain oxygens consistently engaged the lysine ε-amino group, while in EpmA they could additionally adopt a bidentate clamp on the substrate β-amino group, a structural fingerprint of EpmA-type β-recognition. Within either enzyme, no static pocket feature separated productive from non-productive substrate poses, consistent with ligand-binding geometry being only one of several catalytic stages.

Because cofolding models can introduce local pose distortions that bias internal-coordinate strain, all complexes were energy-minimized under the AMBER14 force field via the YASARA Structure suite prior to strain evaluation. Criterion 5 then combines four sub-scores: three scored (AS identity, strain hedge, bid-carrier) that contribute to the composite and decompose into the curves in the criterion-5 panel of main text Fig. 1, plus pose-in-pocket, a sanity-check diagnostic reported per cell but not folded into the composite.

**AS identity** captures active-site sequence retention over 11 disruptable EpmA active-site residues curated from the literature (A298 excluded as the engineered specificity handle). Each position is tiered as *essential*, *heavy*, or *mild* with a corresponding bin weight; a chimera position scores 0 if identical to EpmA, a fractional negative score for conservative substitution (same physicochemical class), or the full negative bin weight for non-conservative substitution. The per-chimera AS-identity score is the sum over the 11 positions.

**Pose-in-pocket** verifies that the cofolding model placed the *(R)*-β-Lys-AMP intermediate inside a plausible binding pocket. A model is considered pocketed if its ligand is contacted by at least 8 protein residues (Cα within 8 Å of any ligand heavy atom); the per-chimera score is the ensemble fraction of pocketed models. Every chimera in the current cohort scores 1.0, so the diagnostic has no exclusionary effect here; it is retained as part of the criterion-5 framework for prospective application to extended chimera sets where ligand mispocketing is more common.

**Strain hedge** captures geometric ligand fit. After AMBER14 minimisation, the bound *(R)*-β-Lys-AMP intermediate is scored against PDB-pooled reference distributions of bond lengths and valence angles, yielding a per-model root-mean-square z-score (rms_z). The per-chimera strain-hedge score is a three-way *hedge* over the n = 10 ensemble combining the pass-rate (fraction of models with rms_z below a calibrated threshold), the ensemble median rms_z, and the ensemble best rms_z, in the same spirit as the criterion-3 hedge.

**Bid-carrier** captures the catalytic-Glu engagement specifically. For each model, the two Glu sidechain oxygens (OE1, OE2) are checked against the two ligand amino groups: the *carrier-N* (the β-amino, adjacent to the activated carboxylate carbon and chemically active in a β-substrate) and the *ε-amino* (NZ, the distal sidechain amine). The pose is classified as **carrier-bidentate** if both oxygens are within 4.5 Å of the carrier-N, the EpmA-type β-engagement signature; **NZ-bidentate** if both oxygens engage the ε-amino instead, the LysS-type α-engagement signature; **bridging** if each oxygen engages a different nitrogen; or **none** if neither is reached. The per-chimera bid-carrier score is the ensemble fraction of carrier-bidentate models.

The three scored sub-scores (z-scored to the cohort, shown at ×⅓ magnitude in the figure so their visual sum equals the composite curve) are combined into the criterion-5 composite as their unweighted mean, evaluated on the (R)-β-Lys-AMP-bound configuration. Separately, the A298 ↔ A298G difference in the bid-carrier sub-score (Δbid-carrier = bid-carrier(A298) − bid-carrier(A298G)) is reported as a *specificity-gate preservation* signal, since a faithful chimera should show diminished EpmA-type β-engagement in the mutant context. As an internal control, EpmA + (R)-β-Lys-AMP anchors the top end of the bid-carrier axis, while LysS + (R)-β-Lys-AMP lands in the NZ-bidentate class.

#### S7. Modelling envelope: split-point range and ligand configurations

Because the engineering objective is a chimera that retains EpmA β-lysine chemistry while gaining LysS-derived tRNA-binding capacity, very high split points are *a priori* unlikely to be productive: by the time the EpmA catalytic core has been progressively replaced by its LysS equivalents, donor specificity will also have become LysS-like. We therefore restricted the sweep to the alignment range up to the column corresponding to EpmA Arg100 (≡ LysS Arg263 in the hybrid map), an essential, class-II-conserved anchor of the ligand phosphate group. This yields 76 unique chimera sequences spanning sp0 to sp100 after sequence-duplicate collapse.

For each chimera, we modelled six ligand configurations chosen to jointly probe folding, ABD engagement, tRNA accommodation, catalytic-fold retention, and β-lysine specificity: (i) apo; (ii) the cognate holo complex with *(R)*-β-Lys-AMP; (iii) the holo complex with (S)-α-Lys-AMP, the canonical (*S*)-α-amino-acid isomer that is the LysS cognate and the preferred substrate of the EpmA A298G mutant; (iv) the holo complex with (*R*)-α-Lys-AMP, an α-stereoisomer outgroup chosen because no (*R*)-β-Lys-AMP-bound class-II structure exists and α-substrate kinetic data are abundant; (v) the chimera with *E. coli* $tRNA^{Lys}$ alone (no donor); and (vi) the chimera with $tRNA^{Lys}$ and (R)-α-Lys-AMP co-bound. All six configurations were predicted in both active-site backgrounds, A298 wild-type and A298G mutant, yielding 12 (variant, configuration) cells per split point and 912 unique complexes across the 76 chimeras of the sweep.

#### S8. Sampling reproducibility and ensemble sizing

To distinguish real structural signal from Boltz seed noise, we audited per-submetric reproducibility from independent duplicate n = 10 Boltz runs of selected chimeras. For each (criterion, metric, configuration) submetric cell, we z-scored each ensemble mean against the cohort distribution of per-chimera ensemble means and plotted z(orig) vs. z(dup). Points within ±1 cohort-SD of the y = x diagonal contribute identically to the composite ranking under either run, i.e. the duplicate would assign the same cohort-relative score. Across the audit, most cells fell well inside this band (Figure S4); the criterion-1 panel showed 5 of 80 cells outside it (a 6 % per-cell outlier rate, considered acceptable at the cohort-rank scale at which the composite operates), but criterion 3, which is fed by a single ligand configuration (the tRNA-only complex) feeding four submetrics, showed 3 of 8 cells outside the band, an outlier rate too high to leave uncorrected (Figure S5). We therefore used n = 10 ensembles for each of the criteria configurations 1, 2, 4, and 5, and increased sampling to n = 40 for the tRNA-containing configurations, bringing the total to 13,680 Boltz structures.

#### S9. Per-criterion structural trends across the sweep

The five criteria each revealed characteristic and largely independent patterns across the split-point range.

**Criterion 1 (dimer integrity).** Confidence metrics rose gradually from sp0 to sp16, then plateaued, with nearly all chimeras in the plateau region exhibiting high-confidence predictions (Figure S6). The tRNA-containing configurations were a partial exception, consistent with the comparatively lower reliability of cofolding models for protein–nucleic-acid complexes.

**Criterion 2 (ABD orientation).** The ABD of sp0–sp9 chimeras regularly adopted dramatically different orientations compared to the LysS crystal reference, in many cases shifting away by several Å or rotating into an alternative pose (Figure S7). From sp10 onward, two well-behaved tiers emerged, sps 10–46 and sps 92–100 with ABD Cα-RMSDs around 1 Å, and sps 47–91 just below 1 Å. As expected of structure prediction models in general, the A298 ↔ A298G point mutation produced no detectable difference in criteria 1 or 2.

**Criterion 3 (tRNA accommodation).** Across the sp range, the $tRNA^{Lys}$ placement engaged the ABD and active site at roughly comparable rates, but past sp51 the exact pose deteriorated moderately, as detected by both the CCA-arm angle and the tRNA rigid-body dihedral (Figure S8). The A298 and A298G variants differed for some chimeras without a clear sp-monotonic pattern.

**Criterion 4 (catalytic-fold retention).** As expected, the EpmA-likeness of the catalytic-domain Cα geometry decreased monotonically with sp, with a notable acceleration at the sp46 → sp47 transition (Figure S9). Per-residue analysis identified two hotspots driving the drift on top of the slow background: the N-terminus and the active-site loop that adopts different conformations in LysS and EpmA.

**Criterion 5 (active-site competence).** Criterion 5 also disfavoured later chimeras. The score was driven down first by penalties for active-site residue exchanges in the loop around residue 50 and at position 76, then by deterioration of the post-minimization ligand strain metric past sp80, and by gradual loss of EpmA-like catalytic-Glu engagement after sp72 (Figures S10, S11).

#### S10. Composite ranking, tier assignment, and lead chimera selection

To aggregate the five criteria into a single ranking, criteria 1 and 2 were applied as fail/pass gates, excluding chimeras whose fold integrity or ABD orientation scored catastrophic. Gate-passing chimeras were z-scored on all five criteria and summed into a composite score per (split point, variant) (Figure S12). The criterion-5 specificity-gate score (the A298 ↔ A298G Glu-engagement difference) was reported separately. To unify the assessment into a single decision-grade metric, all chimeras were assigned to one of seven performance tiers via a weighted combination of the composite score and the specificity-gate signal, with greater weight on the composite, since the primary engineering goal was unlocking *(R)*-β-Lys activity and α-specificity recovery in the mutant was a secondary concern.

The resulting tier landscape was internally coherent. The earliest chimeras, sp0–sp10, failed the criterion 1–2 gate outright due to catastrophic folds and/or ABD misorientation; sps 11–13 cleared the gate but still ranked low for the same reasons, confirming the *a priori* assertion that the drifted EpmA N-terminus is unsuited to directly serve as a fusion point for ABD grafting. The best-performing range of chimeras then ensued from sp15 to sp49, where the variant-mean z-score remained positive throughout. After sp50, performance dropped again, with a local minimum at sp53 and rapid deterioration past sp90. The decline in the sp50–sp90 range was initially dominated by deteriorating tRNA accommodation, while catalytic-domain retention and active-site competence became the dominant penalty terms in the highest-chimeras, confirming the sweet-spot hypothesis that motivated the sp≤100 modeling envelope (Figures 1, S12).

Nine top-ranking (sp, variant) cells fell in the sp24–sp49 range, corresponding to eight unique chimera sequences. Within this top set, no single chimera dominated all assessment axes simultaneously. sp45, for instance, ranked highest on the A298 variant, but its mutant ranked rather low, and sp49 showed the opposite pattern; sp2, while having the lowest variant-mean z-score among the winners, outranked the others in specificity-gate preservation. As a compromise that combined an excellent A298 composite score with good preservation of the specificity-gate signal, we selected sp43 as our lead chimera. We additionally chose it because it is one of the most progressed designs still safely upstream of the active-site loop that begins to be substituted from sp49 onward. Within the sp15–sp38 range, we observed a moderate oscillation in the combined z-score between adjacent split points, which we interpret as evidence of the structural scoring's sensitivity to discrete sequence-level changes between neighboring chimeras.

### Experimental Section

#### Materials and chemicals

Chemicals and reagents were obtained from Sigma-Aldrich (Taufkirchen, Germany), Carl Roth (Karlsruhe, Germany), or Thermo Fisher Scientific (Waltham, MA, USA) at the highest available purity unless otherwise stated. ATP disodium salt and deoxyribonuclease I (DNase I, from bovine pancreas) were purchased from Sigma-Aldrich. RNase-free DNase I was from Thermo Fisher Scientific. γ-[$^{32}$P]-ATP was supplied by Hartmann Analytic (Braunschweig, Germany). Ni-NTA agarose was from Qiagen (Hilden, Germany). T7 RNA polymerase and ribonucleotide triphosphates (rNTPs) were from New England Biolabs (NEB, Ipswich, MA, USA). *N*-α-Boc-lysine, glyoxylic acid monohydrate, and palladium on charcoal were obtained from Sigma-Aldrich. *(R)*-β-lysine, $N^{ε}$-acetyl-(*S*)-α-lysine, $N^{ε}$-succinyl-(*S*)-α-lysine, and biocytin were from Bachem (Bubendorf, Switzerland) or Sigma-Aldrich. $N^{ε}$-carboxyethyl-(*S*)-α-lysine (CEL) was synthesised in-house following the procedure of Hellwig *et al*.[1] Antibiotics (ampicillin sodium salt, kanamycin sulfate, chloramphenicol, tetracycline hydrochloride, spectinomycin), L-(+)-arabinose, and isopropyl β-D-1-thiogalactopyranoside (IPTG) were from Carl Roth. DOWEX 50W-X8 cation-exchange resin (100–200 mesh) was from Sigma-Aldrich.

##### Synthesis of $N^{ε}$-carboxymethyl-(*S*)-α-lysine (CML)

*N*-α-Boc-lysine (2.06 g, 8.4 mmol) and glyoxylic acid monohydrate (970 mg, 10.5 mmol) were dissolved in 60 mL of water, and the pH was adjusted to 8.75 with diluted sodium hydroxide solution. Palladium on charcoal (100 mg) was added, and the mixture was stirred for 72 h at room temperature under a hydrogen atmosphere. After filtration, 60 mL of 6 M HCl was added, and the mixture was stirred at room temperature for 3 h. Hydrochloric acid was then evaporated *in vacuo*, and the residue was dissolved in 50 mL of 0.01 M HCl. The solution was applied to a column (25 × 2.5 cm) filled with the strongly acidic cation-exchange resin DOWEX 50W-X8 (100–200 mesh), previously equilibrated with 300 mL of 6 M HCl, 300 mL of water, and 100 mL of 0.01 M HCl. Elution was performed first with 50 mL of 0.01 M HCl, then with 700 mL of 1.5 M HCl. Fractions (6 mL) were collected with a fraction collector (Model 2110, Bio-Rad Laboratories, Feldkirchen, Germany). Target compounds were detected by spotting 1 µL of each fraction on TLC plates and spraying with a solution of 0.1 % ninhydrin in ethanol, followed by heating to 80 °C. CML was found to elute between 450 and 700 mL of 1.5 M HCl. Positive fractions were pooled, filtered, and evaporated to dryness *in vacuo*. After lyophilization, CML dihydrochloride was obtained as a white powder and stored at –18 °C.

**Analytical data:** mass spectrometry, $[M + H]^+$ *m/z* = 205.0; elemental analysis, calcd. for $C_8H_{16}N_2O_4$ × 2 HCl (MW = 277.1): C, 34.67; H, 6.55; N, 10.11. Found: C, 33.80; H, 6.44; N, 9.90; content 98 % based on nitrogen. Yield: 1.18 g (49.7 % from lysine).

##### Bacterial strains and growth conditions

*Escherichia coli* strains were routinely grown aerobically in Luria-Bertani (LB) medium [1% (w/v) NaCl, 1% (w/v) tryptone, 0.5% (w/v) yeast extract, pH 7.0] at 37 °C. Antibiotics were used at the following final concentrations: 100 µg mL$^{-1}$ ampicillin sodium salt, 50 µg mL$^{-1}$ kanamycin sulfate,

30 μg mL$^{-1}$ chloramphenicol, 15 μg mL$^{-1}$ tetracycline hydrochloride, and 50 μg mL$^{-1}$ spectinomycin. Growth curves were recorded using a Spark 20M multimode microplate reader (Tecan, Männedorf, Switzerland) at 37 °C; optical density at 600 nm ($OD_{600}$) was measured every 20 min. Strains used in this work are listed in Table S1.

Plasmid construction

For CRISPR interference (CRISPRi) targeting of the chromosomal *lysS* gene, the psgRNA backbone[2] was used. A 20-nt spacer (5′-CATTAAAGATCTGCGCACCGG-3′) located 115 nucleotides upstream of the *lysS* +1 site was inserted into psgRNA by Gibson assembly (Table S2).

For chromosomal integration of chEpmA with concomitant deletion of *lysS*, the CRISPR/Cas9-based two-plasmid system of Jiang *et al.* was used[3]. A 20-nt protospacer specific to the *lysS* coding sequence was cloned into the pTarget plasmid (Table S2). A homologous recombination donor was constructed by flanking the chEpmA-α coding sequence with 500 bp arms corresponding to the upstream and downstream genomic regions of *lysS* (Table S2). All plasmids used in this work are listed in Table S3.

**Genome editing**

Competent *E. coli* BW25113 Δ*lysU* cells previously transformed with pCas9[1] were cultured overnight in LB supplemented with 50 μg mL$^{-1}$ kanamycin at 37 °C with shaking. The culture was diluted 1:100 into LB containing kanamycin (50 μg mL$^{-1}$) and L-(+)-arabinose (10 mM final) to induce λ-Red recombinase expression. When the $OD_{600}$ reached 0.5, 1 mL of cells was harvested by centrifugation and washed twice with sterile 10 % glycerol. The washed cells were resuspended in 200 μL of 10 % glycerol and mixed with 50 ng of pTarget plasmid and 400 ng of the homologous recombination fragment. Electroporation was performed in a 2-mm Gene Pulser cuvette (Bio-Rad) at 2.5 kV, and cells were immediately resuspended in 1 mL of ice-cold LB. After 1 h of recovery at 30 °C, cells were plated on LB agar containing kanamycin (50 μg mL$^{-1}$) and spectinomycin (50 μg mL$^{-1}$) and incubated overnight at 30 °C. Transformants were verified by colony PCR and Sanger sequencing.

To cure the pTarget plasmid, a colony harboring both pCas9 and pTarget was inoculated into 2 mL of LB containing kanamycin (50 μg mL$^{-1}$) and IPTG (0.5 mM) and incubated for 8–16 h at 30 °C. The culture was then diluted and spread onto LB plates containing kanamycin (50 μg mL$^{-1}$). Colonies were screened for sensitivity to spectinomycin (50 μg mL$^{-1}$); spectinomycin-sensitive isolates were considered cured of pTarget. To eliminate pCas9, cured colonies were grown overnight at 37 °C without antibiotic selection.

**Protein production and purification**

All recombinant proteins were produced in *E. coli* LMG194 carrying the corresponding pBAD33-based expression vectors (pBAD33_LysS, pBAD33_chEpmA-α, pBAD33_EpmA; Table S3). Expression was induced during exponential growth by adding 0.2% (w/v) L-(+)-arabinose, and cultures were incubated overnight at 18 °C. Cells were harvested by centrifugation and resuspended in HEPES lysis buffer [50 mM HEPES, 100 mM NaCl, 50 mM KCl, 10 mM $MgCl_2$, 5 % (w/v) glycerol, pH 7.0] supplemented with DNase I, then incubated for 30 min at 4 °C. Cells were disrupted using a continuous-flow cell disruptor (Constant Systems Ltd., Daventry, UK) operating at 1.35 kbar. Lysates were clarified by ultracentrifugation at ~235,000 × *g* for 1.5 h at 4 °C. $His_6$-tagged proteins were purified on Ni-NTA agarose (Qiagen, Hilden, Germany) following the manufacturer's instructions: after washing with HEPES buffer containing 20 mM imidazole, target proteins were eluted with HEPES buffer containing 400 mM imidazole. Pooled fractions were dialyzed overnight at 4 °C against imidazole-free HEPES buffer. Protein purity was assessed by SDS-PAGE, and concentrations were determined spectrophotometrically (NanoDrop, Thermo Fisher Scientific). Aliquots were flash-frozen in liquid nitrogen and stored at –80 °C.

##### *In vitro* tRNA synthesis

*E. coli* $tRNA^{Lys}$ was produced by run-off *in vitro* transcription. The template was a synthetic double-stranded DNA fragment in which the T7 RNA polymerase promoter was placed immediately upstream of the $tRNA^{Lys}$ coding sequence (Table S4); a single-stranded primer complementary to the T7 promoter was used to prime the reaction (Table S4). Transcription reactions (160 µL final volume) were assembled in T7 reaction buffer (16 µL of 10× stock) with template (0.5 µg), primer (32 µL of 20 µM stock), rNTPs (0.5 mM each final concentration; 32 µL), and T7 RNA polymerase (8 µL; NEB) in nuclease-free $H_2O$. Reactions were incubated at 37 °C for 16 h. Subsequently, 2 µL of RNase-free DNase I (Thermo Fisher Scientific) was added, and the mixture was incubated for an additional 2 h at 37 °C to digest the DNA template. The tRNA was purified by ethanol precipitation. Each transcription reaction was supplemented with 200 µL of nuclease-free $H_2O$ and 30 µL of 3 M sterile sodium acetate, then mixed with 2.5 volumes of 99 % ethanol and incubated at –20 °C for 1 h. Samples were centrifuged at 10,000 × *g* for 30 min at 4 °C; the supernatant was discarded, and the pellet was washed with 500 µL of 70 % ethanol, centrifuged at 10,000 × *g* for 15 min at 4 °C, air-dried, and resuspended in 50–100 µL of nuclease-free $H_2O$. tRNA concentration and integrity were assessed by UV absorbance.

##### Aminoacylation activity assay ([$^{32}$P]-$PP_i$ release)

tRNA charging activity was monitored by quantifying the release of [$^{32}$P]-pyrophosphate ([$^{32}$P]-Aminoacylation activity assay. tRNA charging activity was monitored by quantifying the release of [$^{32}$P]-pyrophosphate ([$^{32}$P]-PPi) from γ-[32P]-ATP, exploiting the fact that organic (nucleotide-bound) phosphate adsorbs to activated charcoal whereas free inorganic phosphate does not. A quenching mix (7% perchloric acid, 10 mM sodium pyrophosphate, 3% w/v activated charcoal) was aliquoted (250 µL per sample) into 1.5 mL reaction tubes. Each aminoacylation reaction (60 µL final volume) was assembled by combining 20 µL of a 3×-concentrated enzyme-free mix with 40 µL of a 1.5×-concentrated substrate mix, yielding final concentrations of 150 mM HEPES pH 7.5, 10 mM MgCl2, 3 mM ATP (including a trace amount of γ-[$^{32}$P]-ATP, 0.01 µM), and 4 U $mL^{-1}$ inorganic pyrophosphatase. Unless stated otherwise, reactions contained the amino acid substrate at a saturating concentration (1 mM), $tRNA^{Lys}$ at 15 µM, and 1 µM enzyme; $tRNA^{Lys}$

concentrations were varied from 0 to 50 µM for Michaelis-Menten analysis. Reactions were incubated at 37 °C for times ranging from 30 s to 16 h, as specified in the corresponding figure legends. At each time point, a 10 µL aliquot was withdrawn into the quenching mix, vortexed vigorously, and, once all samples had been collected, the suspensions were centrifuged for 5 min at 10,000 × g. Aliquots (50 µL) of the supernatant were added to 5 mL scintillation cocktail, and the released [$^{32}$P]-phosphate (following pyrophosphatase-mediated hydrolysis of [$^{32}$P]-PPi) was quantified by liquid scintillation counting. All values were baseline-corrected by subtracting the corresponding tRNA-only control. For Michaelis-Menten analysis, initial velocities were determined from the amount of aminoacylated product formed after a fixed 10-min reaction time at each $tRNA^{Lys}$ concentration, assuming that product formation remained within the initial-rate (linear) regime over this interval; initial velocity ($v_0$) was calculated as [product]/reaction time. Steady-state kinetic parameters ($K_M$ and $k_{cat}$) were obtained by fitting $v_0$ versus $tRNA^{Lys}$ concentration to the Michaelis-Menten equation by weighted nonlinear regression. All values are reported as mean ± SEM from three independent replicates.

#### Computational design of chEpmA chimeras

*Inventory of ground-truth structures*

Crystal structures of EpmA, LysS, and class-II aminoacyl-tRNA-synthetase homologs were retrieved from the RCSB Protein Data Bank (PDB) using two complementary searches. A structural homology search was performed with FoldSeek [4] against the public PDB database (pdb100 target set), seeded with the *E. coli* EpmA crystal (PDB 3A5Z, the EpmA–EF-P complex, 323-residue full-length chain) in 3diaa mode and using default e-value filtering. In parallel, a sequence-based search was executed via the RCSB Data API: a GraphQL query was sent over HTTPS POST (Python urllib.request, endpoint https://data.rcsb.org/graphql) for all PDB entries matching the EpmA/LysS EC numbers and gene names with ≥ 30 % sequence identity to the E. coli sequences (UniProt P0A8N7 for EpmA, P0A8N3 for LysS); this sequence-identity search also surfaced the LysS family hits. The two result sets were merged, deduplicated, and manually screened for false positives (e.g. entries returned by the EC-number match that proved to be unrelated enzymes upon inspection), yielding a master reference table. From this table, the EpmA + (S)-α-Lys-AMP analogue complex was used as the EpmA non-cognate-substrate pose reference, the LysU + (S)-α-Lys-AMP complex was used as the productive-cognate pose reference for the LysS lineage (LysU is 89 % identical to *E. coli* LysS), and the human cytosolic LysS–tRNA complex (PDB 9DPL) was used as the tRNA-bound reference at this fold, as no bacterial LysS–tRNA complex has been crystallized to date.

*Structure-guided EpmA–LysS alignment*

The EpmA + *(R)*-β-Lys-AMP (PDB 3A5Y) and the LysS apo (PDB 1BBU) catalytic-domain crystals were retrieved from RCSB and loaded into a YASARA Structure session via the YASARA Python API. A structural alignment of the two catalytic domains was computed with SHEBA[5], invoked through the YASARA AlignObj function with method='SHEBA'. The alignment operates on Cα atoms only and yields a list of structurally equivalent residue pairs at sub-Ångström geometric proximity, together with the underlying Cα superposition transform. The SHEBA-anchored residue pairs define the canonical structurally equivalent positions; the resulting residue-equivalence map covers the catalytic-domain core but leaves unanchored (i) residues unresolved in one or both crystals and (ii) pairs that lie too far apart in 3D for SHEBA to call structurally equivalent.

The unanchored alignment columns were filled with a pairwise BLOSUM62[6] sequence alignment computed in Python with the Biopython[7] Bio.Align.PairwiseAligner class using substitution_matrices.load('BLOSUM62'). For all internal (non-terminal) regions, gap-open and gap-extend scores were set to −11 and −1, respectively. For the N- and C-terminal regions that extend beyond the SHEBA anchors, a semiglobal aligner with stricter internal gap penalties (open/extend = −30/−2) and free-end penalties on the alignment-extension side was used to discourage internal gap-opening while permitting flush end alignment.

The composite output of the SHEBA structural-anchor table and the BLOSUM62 fill is a single residue-equivalence map (one entry per alignment column, with the EpmA residue index and the LysS residue index for each anchored position) that is the canonical input for chimera sequence generation and for all per-residue structural measurements that compare a chimera to either parent.

*Chimera sequence generation*

Chimera sequences were generated programmatically from the residue-equivalence map by the make_chimera_seqs.py script. At a given split point n, the chimera amino-acid sequence is constructed as the concatenation of (i) the *E. coli* LysS anticodon-binding domain (residues 1–167 of the full-length LysS sequence), (ii) the LysS residues mapped to alignment columns 0 through n − 1, and (iii) the EpmA residues mapped to alignment columns n through the C-terminus, with gap characters from the aligned strings removed. The procedure naturally handles indels in the alignment: at an EpmA-deletion column moving from spn to spn + 1 adds one LysS residue while the EpmA portion is unchanged, and at an EpmA-insertion column the LysS portion is unchanged while EpmA loses one residue. The script emits one FASTA file per (split point, active-site variant) pair, where the variant is either the wild-type A298 background or the A298G point mutant introduced by direct sequence substitution at the EpmA-mapped active-site position. Adjacent split points yielding byte-identical chimera sequences (i.e., spanning runs of residue identity between EpmA and LysS) were collapsed to a single canonical split point downstream by md5 comparison of the generated sequences.

*Ligand model preparation*

The six ligand configurations modeled in the sweep, apo, *(R)*-β-Lys-AMP, (S)-α-Lys-AMP, (R)-α-Lys-AMP, *E. coli* tRNA-Lys, and tRNA-Lys co-bound with (R)-α-Lys-AMP, were prepared as canonical SMILES strings (small molecules) or as residue-level RNA sequences (tRNA), defined in make_physiological_smiles.py. Each Lys-AMP-class SMILES carries explicit (R)/(*S*) stereo annotations at the α-carbon, explicit α- vs β-backbone connectivity at the lysine carboxylate, and protonation states consistent with the physiological pH 7.4 (zwitterionic lysine sidechain protonated, α-phosphate doubly deprotonated, β-phosphate group of AMP doubly deprotonated). The stereochemistry of each ligand was verified before use by loading the SMILES into RDKit [8], calling Chem.AssignStereochemistry(mol, cleanIt=True, force=True), and confirming that the assigned CIP descriptor (_CIPCode atomic property) at the α-carbon matched the intended *(R)* or (*S*) label. The *E. coli* tRNA-Lys sequence was used in its unmodified primary sequence. Identical ligand inputs were used across all chimera × variant cells for each ligand configuration.

*Structure prediction by Boltz-2 cofolding*

All protein–ligand and protein–nucleic-acid complexes were predicted with the Boltz-2 cofolding model [9] via its command-line interface (boltz predict). For each (chimera, active-site variant, ligand configuration) cell, a Boltz-2 input YAML was generated programmatically, referencing (i) the chimera FASTA produced by the chimera-generation step, (ii) the appropriate ligand SMILES (or tRNA-Lys sequence; or both, for the co-bound configuration), and (iii) the multiple-sequence alignment (MSA) for the chimera protein chain.

For each chimera, the MSA was generated once with the ColabFold MMseqs2 server [10] (Boltz-2 --use_msa_server flag), then cached as an .a3m file and reused across the two active-site variants (A298 / A298G) of the same chimera, since the single-residue mutation does not warrant re-fetching the MSA. Each prediction was run with --recycling_steps 10, --diffusion_samples 10, --sampling_steps 200, and --use_potentials. The --diffusion_samples 10 setting yields an ensemble of 10 independent stochastic samples (mdls) per (chimera, variant, configuration) cell for the four single-ligand configurations (apo, *(R)*-β-Lys-AMP, (*S*)-α-Lys-AMP, (R)-α-Lys-AMP); for the two tRNA-containing configurations, four independent Boltz-2 runs of --diffusion_samples 10 were aggregated to yield an n = 40 ensemble per cell (see *Ensemble sampling and reproducibility audit*, below). All predictions were executed under SLURM scheduling on the Snellius (SURF) and Habrok (University of Groningen) GPU clusters, with each job requesting 1 × GPU (NVIDIA H100 on Snellius; NVIDIA A100 or RTX Pro 6000 on Habrok), 32 GB RAM, 7 CPU threads, and an 8 h walltime.

*Ensemble sampling and reproducibility audit*

To set the per-cell ensemble size, an independent duplicate Boltz-2 run (also --diffusion_samples 10) was generated for a representative subset of chimeras. For every (criterion, metric, ligand configuration) submetric cell, the per-chimera ensemble mean of each duplicate was z-scored against the cohort distribution of per-chimera ensemble means (chimera-only cohort, wild-type references excluded; cf. *Composite scoring*, below). Reproducibility of a submetric cell was scored as the fraction of chimeras whose two duplicate z-scores lay within ±1 cohort standard deviation of the y = x diagonal, i.e. the band within which a duplicate run assigns the same cohort-relative score and contributes identically to the composite ranking. For submetrics in criteria 1, 2, 4, and 5, this fraction was sufficient at n = 10. For criterion 3 — the only criterion fed by a single ligand configuration (tRNA-Lys) and a single per-cell metric count of four — the per-cell reproducibility at n = 10 was below the acceptable threshold. The ensemble size for the tRNA-containing configurations was therefore set to n = 40, pooled across four independent Boltz-2 --diffusion_samples 10 runs of the same input YAML (differing only in random seed).

*Per-criterion structural measurement*

All per-criterion measurement scripts read the Boltz-2 prediction outputs (CIF + confidence JSON) and emit a per-(chimera, variant, configuration, model) row. YASARA-dependent steps were performed by importing the yasara Python module (from yasara import *) shipped with the YASARA Structure suite, which exposes the YASARA command set as Python functions and is the standard interface for scripted YASARA analysis.

**Criterion 1: structural integrity**. For each predicted model, the Boltz-2 confidence outputs were parsed to extract per-residue pLDDT (aggregated as the catalytic-domain mean mean_plddt_cat and the ABD mean mean_plddt_abd), the chain-level pTM (chain_0_ptm), and the homodimer interface pTM (protein_iptm). A predicted dimer Cα-RMSD between the two protein chains of the homodimer (dimer_rmsd_AB) was computed via YASARA Sup; a total steric-clash count (clash_tot) was derived from YASARA's ListAtomVdwClash over the dimer; and the longest contiguous stretch of catalytic-domain residues with pLDDT < 70 (longest_low_cat) was enumerated to flag locally disordered regions that the catalytic-domain mean pLDDT averages over. These seven quantities are aggregated per chimera by averaging over the n = 10 ensemble and (for the *positive* components) across the canonical configuration set, then z-scored against the chimera cohort and combined into a single integrity score by

integrity_score = 2 · z(mean_plddt_cat) + 1 · z(mean_plddt_abd) + 1 · z(chain_0_ptm) + 1 · z(protein_iptm) − 1 · pen(dimer_rmsd_AB, 1.0 Å) − 1 · pen(clash_tot, p95_EpmA-WT) − 0.5 · pen(longest_low_cat, 5 residues)

where pen(x, t) = max(0, (x − t)) / σ(x) is the threshold-excess penalty, normalised by the chimera-cohort standard deviation of x. Computation is implemented in score_structural_integrity.py.

**Criterion 2: inter-domain orientation**. Since no full-length *E. coli* LysS crystal containing the ABD has been solved, the reference for ABD orientation is the human cytosolic LysS chain A from the LysS–tRNA complex (PDB 9DPL), which carries the full ABD in a physiological orientation relative to its catalytic domain. Each predicted chimera was SHEBA-aligned to 9DPL chain A in YASARA, and the Cα-RMSD of the chimera's ABD residues was then computed against 9DPL over a fixed 132-residue chimera-ABD-to-9DPL-ABD correspondence map (abd_resnum_map_9dpl.json). The correspondence map was derived once, by SHEBA-aligning the *E. coli* LysS-WT apo Boltz-2 model 1 onto 9DPL chain A and harvesting the pairs called structurally equivalent; using this fixed pair set (rather than running SHEBA per chimera) ensures that a mis-placed ABD region still enters the RMSD with its native pairs rather than being silently re-anchored at a spurious local minimum. The script measure_global_geometry_yasara.py emits one row per (chimera, variant, configuration, model). The criterion-2 composite per (split point, variant), the geometry score, is the pooled mean of per-model ABD Cα-RMSDs across the four apo-like ligand configurations (apo, (*R*)-β-LysAMP, (*R*)-α-LysAMP, (*S*)-α-LysAMP), plus tRNA-bound models that pass an upstream per-model tRNA-dock-coherence gate (trna_well_docked, requiring criterion-3 productivity in that model); the latter pre-filter prevents tRNA-bound models with catastrophic dock geometry from contaminating the resting-state-orientation signal, while allowing well-docked tRNA frames to contribute. The metric is sign-flipped (higher RMSD = worse) when combined into the composite.

**Criterion 3: tRNA accommodation**. Each predicted chimera–tRNA complex was aligned over the chimera catalytic-domain Cα atoms to the human LysS–tRNA reference (PDB 9DPL), and four geometric quantities were measured via YASARA primitives: (i) the distance d_acc from the tRNA CCA-end (3′-acceptor terminus) to the chimera's active-site Glu; (ii) the distance d_antic_patch from the anticodon-loop centroid to a fixed ABD-anchor patch on the chimera; (iii) the cosine cos_arm_angle of the angle between the tRNA acceptor-arm vector and the anticodon-arm vector, capturing the bend at the L-shape elbow (the "kink"); and (iv) the cosine cos_plane_dihedral of the dihedral about the elbow axis, capturing rotation of the two arms relative to each other. The four quantities are combined into a single per-model productivity score by summing four penalty terms — sigmoid penalties on d_acc above 9 Å and on d_antic_patch above 6 Å, a linear penalty on cos_arm_angle below the wild-type-reference mean, and a quadratic penalty on cos_plane_dihedral outside the wild-type-reference window — and a model is considered passing if its productivity score exceeds −10, a hard threshold chosen so that the wild-type chimera-reference complexes (whose productivity score is ≈ 0) pass comfortably while the catastrophic regime is clearly excluded. Per-model values are emitted by measure_trna_accommodation_yasara.py. Aggregation across the n = 40 ensemble of each (chimera, variant) cell produces three summary statistics — the best (most productivity-score-consistent) model, the median model, and the pass-rate (fraction of models exceeding −10) — which are each z-scored against the chimera cohort and combined into a single criterion-3 *hedge* score = mean(z_best, z_median, z_pass-rate) so that outlier-best, typical-case, and robustness signals are weighted equally.

**Criterion 4: catalytic-fold retention**. For each predicted chimera, the catalytic-domain Cα atoms were independently superposed onto the EpmA catalytic-domain crystal (PDB 3A5Y, the EpmA + (R)-β-Lys-AMP complex) and onto the LysS catalytic-domain crystal (PDB 1E1T as the primary reference, with 1BBU as a secondary cross-check), each in a separate YASARA Sup operation. From each superposition, the per-residue Cα–Cα distance between the chimera and the corresponding parent residue was tabulated, yielding two parallel per-residue distance arrays: $d_i$,EpmA and $d_i$,LysS. The chimera's catalytic-fold scalar δ is the residue-summed difference δ = $\Sigma_i$ ($d_i$,LysS − $d_i$,EpmA) over the catalytic-domain residues. A positive δ means the chimera sits geometrically closer to EpmA than to LysS at residue i on average, i.e. the catalytic core has retained EpmA character, while a negative δ flags drift toward the LysS catalytic geometry. δ is therefore monotone in catalytic-core EpmA-likeness, and is sign-flipped before being combined into the composite so that larger composite values continue to indicate better designs. Computation is implemented in measure_catalytic_fold_yasara.py.

**Criterion 5: active-site competence**. Criterion 5 combines four sub-scores, **AS identity**, **pose-in-pocket**, **strain hedge**, and **bid-carrier**, of which the first, third, and fourth contribute to the criterion-5 composite while the second is fulfilled in all models and only reported as sanity-check diagnostic.

**AS identity.** A curated set of 11 disruptable EpmA active-site residues was defined a priori from the literature on EpmA/LysS active-site function (the A298 specificity gate itself is excluded because it is the engineered handle and is reported separately). Each residue position carries a literature-derived disruption-severity tier (essential / heavy / mild) and a corresponding bin weight. For each chimera position, the residue is compared to the EpmA reference and scores 0 if identical to EpmA, the negative bin weight if substituted by a different residue, and a fractional negative score if the substitution is conservative within the same physicochemical class. The chimera's AS-identity score is the sum of per-position contributions over the 11 residues. Computation is implemented in recompute_s1_weighted.py.

**Pose-in-pocket.** For every (R)-β-Lys-AMP-bound model, the bound ligand is checked against a minimum-engagement criterion: the ligand must be contacted by at least 8 protein residues, where contact is defined as having at least one Cα within 8 Å of any ligand heavy atom. The per-chimera pose-in-pocket score is the per-ensemble fraction of models meeting this criterion. The check is conceived as a sanity gate that disqualifies chimeras whose pocket has collapsed or whose ligand has been placed outside any reasonable binding site; in the current chimera cohort every chimera passes it (per-cell score = 1) so the diagnostic is reported alongside but not folded into the composite. Computation is implemented in s2_pose_in_pocket in score_chimeras_criterion5.py.

**Strain hedge.** Each (R)-β-Lys-AMP-bound chimera prediction is energy-minimized in YASARA Structure under the AMBER14 force field [11] using the following recipe, applied to one ensemble member at a time: (i) buffer/cryoprotectant heteroatoms are dropped while preserving any metal-coordinated waters, the structure is oriented, and gap-bridging peptide bonds at unresolved loops are broken so that CleanAll adds ACE/NME caps; (ii) protonation states are assigned at pH 7.4 via YASARA's pKa workflow followed by OptHydAll; (iii) the system is enclosed in a cubic periodic cell with 10 Å padding, parameterised with AMBER14 + SetPar=yes (which auto-types CCD ligands via GAFF2/AM1BCC during CleanAll), set up with PME electrostatics; (iv) the cell is solvated with explicit water and neutralised at 0.9 % (mass fraction) Na/Cl ionic strength via YASARA's Experiment Neutralization; (v) the full system is minimised to convergence at 0.05 kJ/mol/atom. After minimisation, the bound (R)-β-Lys-AMP intermediate is extracted and the internal-coordinate strain of the ligand is scored against PDB-pooled reference distributions of bond lengths and valence angles: each coordinate yields a z-score against its reference distribution, and the per-model strain metric is the root-mean-square of those z-scores (rms_z). The per-chimera strain-hedge score is a *hedge* over the n = 10 ensemble, combining (i) the pass-rate of models whose rms_z falls below a calibrated threshold, (ii) the ensemble median rms_z, and (iii) the ensemble best rms_z, into a single value in [0, 1].

**Bid-carrier.** The catalytic glutamate of the chimera (EpmA Glu116, equivalent to LysS Glu279; the corresponding chimera residue number is resolved per-spvia the alignment map by the helper cat_glu_chim_residue) is checked for *bidentate-on-carrier* engagement of the bound *(R)*-β-Lys-AMP. The "carrier" nitrogen is the lysine amine adjacent to the activated carboxylate carbon — the β-amino group for β-substrates and the α-amino group for α-substrates, identified geometrically per model as the (R)-β-Lys-AMP nitrogen lying closest to the substrate's carboxylate carbon; the ε-amino is the more distal sidechain nitrogen (NZ). For each model, the two Glu sidechain-oxygen-to-nitrogen distances are measured to both nitrogens (OE1→carrier-N, OE2→carrier-N, OE1→NZ, OE2→NZ) and the engagement is classified as: **carrier-bidentate** ("carr") if both OE1 and OE2 are within 4.5 Å of the carrier-N (the EpmA-type β-engagement signature); **NZ-bidentate** ("NZ") if both oxygens are within 4.5 Å of the ε-amino instead (the LysS-type α-engagement signature); **bridging** ("bridge") if one oxygen engages each nitrogen; or **none** if no oxygen is within reach. The chimera's bid-carrier score per configuration is the per-ensemble fraction of models in the "carr" class, computed by s4_bidentate in score_chimeras_criterion5.py.

The criterion-5 composite is the unweighted arithmetic mean of the three scored sub-scores: score_crit5 = (AS_identity + strain_hedge + bid_carrier) / 3, evaluated on the *(R)*-β-Lys-AMP-bound configuration as the catalytically relevant substrate regardless of variant. Separately, the A298 ↔ A298G difference in the bid-carrier sub-score (Δbid-carrier = bid_carrier(A298) − bid_carrier(A298G)) is reported as the *specificity-gate preservation* signal: a faithful chimera should show diminished EpmA-type β-engagement specifically in the mutant context.

*Composite scoring and tier assignment*

The five per-criterion scores were aggregated into a single composite score per (split point, variant) by score_composite.py in three stages.

**Stage 1: gating.** Two fail/pass gates were applied. A chimera fails the criterion-1 gate if its integrity score is below −10 (catastrophic fold confidence); it fails the criterion-2 gate if its ABD Cα-RMSD exceeds 2.5 Å (catastrophic ABD displacement). The two thresholds were calibrated to admit every cohort chimera whose dimer fold and ABD orientation are visually inspectable as a coherent prediction while excluding the catastrophic regime occupied by the earliest split points (sp0–9), where the model returns either disordered or geometrically incoherent dimers; the threshold values were fixed before composite scoring and are hard-coded in score_composite.py. Chimeras failing either gate were excluded from cohort z-scoring and assigned to the bottom performance tier directly.

**Stage 2: cohort z-scoring.** For each of the five criterion scores, the gate-passing chimera cohort distribution (wild-type EpmA and wild-type LysS reference predictions excluded) was used to define a mean and a standard deviation, and each chimera's score was z-transformed against this cohort. The per-criterion z-scores were combined into a single composite by sign-aware summation: composite = z(crit 1) + (−1) · z(crit 2) + z(crit 3 hedge) + (−1) · z(crit 4) + z(crit 5) where the (−1) coefficients flip the sign of distance/penalty metrics (crit 2 ABD-RMSD and crit 4 δ) so that higher composite always corresponds to better. The criterion-5 specificity-gate preservation score was reported separately and not summed into the composite.

**Stage 3: tier assignment.** Chimeras were ranked by composite score and binned with the specificity-gate preservation score (Δbid-carrier) into a seven-bin tier scale ({1.0, 1.5, …, 4.0}) by a weighted average of the per-criterion tier ordinals — tier_avg = (4 · composite_tier + Δbid_carrier_tier) / 5, then snapped to the nearest 0.5, so that the composite carries four times the weight of the specificity-flip signal in the final tier assignment. The 4 : 1 weighting reflects the engineering priority of unlocking *(R)*-β-Lys activity over preserving the A298G α-acceptance switch.

##### *Software, code, and data availability*

All analyses were implemented in Python 3.12 using the following packages with their standard scientific-stack versions: Biopython[7] (sequence alignment), RDKit[8] (ligand SMILES handling, CIP assignment), the YASARA Python module shipped with the YASARA Structure software suite [12–14] (structural alignments, RMSD computation, distance/angle/dihedral primitives, AMBER14 energy minimisation), pandas and numpy (data handling), and matplotlib (figure generation). Structure prediction was performed with Boltz-2 [9] (v2.2.1) via its command-line interface, with the parameters detailed above; MSAs were obtained via the ColabFold MMseqs2 server [10]. Foldseek [4] was used for the PDB structural homology search; the RCSB Data API GraphQL endpoint was queried for the sequence-based PDB search. All analysis scripts, the canonical EpmA–LysS residue-equivalence map, the per-chimera FASTAs, the per-cell Boltz-2 input YAMLs, the per-criterion measurement output tables, and the composite-score table are available at https://github.com/mf-rug/Gallo_2026_chEpmA. Full Boltz-2 prediction outputs (per-mdl CIFs and confidence JSONs) are deposited at https://zenodo.org/records/20990839.

##### *Use of AI coding assistance*

We used Claude (Anthropic; version Opus 4.7; “xhigh” thinking mode) to assist with code generation, data analysis, and visualisation. All code outputs were reviewed, validated, and modified by the authors, who retain full responsibility for accuracy and interpretation.

#### Supplementary Figures

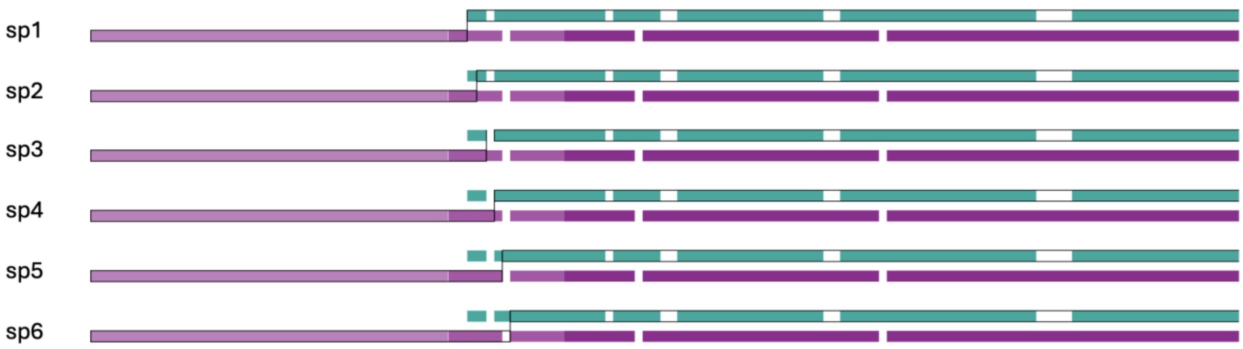


**Figure S1.** Schematic depiction of the split point (sp) sweep. The first chimera's sequence is composed of the N-terminus of LysS (purple) and the complete EpmA sequence (turquoise). Higher-sp chimeras increasingly replace N-terminal EpmA residues with LysS equivalents by moving the sp (black line) C-terminally along the sequence alignment columns. Indels (white) in the alignment cause chimera sequences to bear one of the two portions identical to its neighbouring chimera, while the other portion becomes shorter or longer (sp5 vs sp6 in this example).

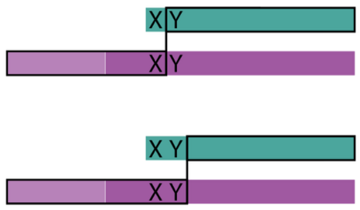


**Figure S2.** Regions with adjacent conserved residues between LysS and EpmA cause chimera neighbours to become sequence identical.

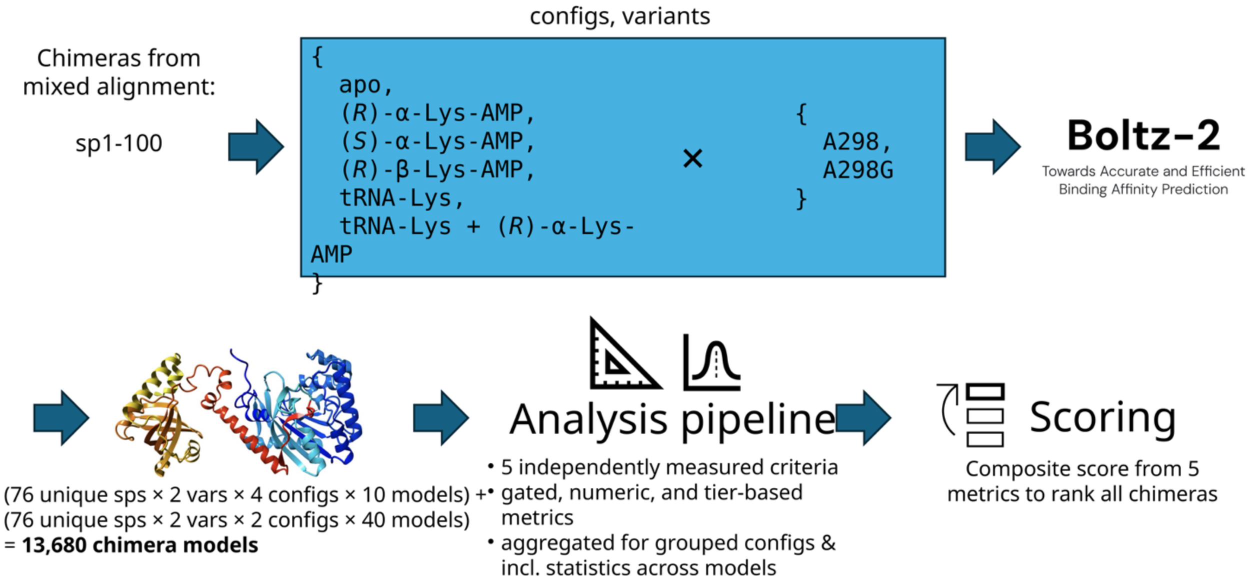


**Figure S3**. Overview of the computational modelling and analysis pipeline.

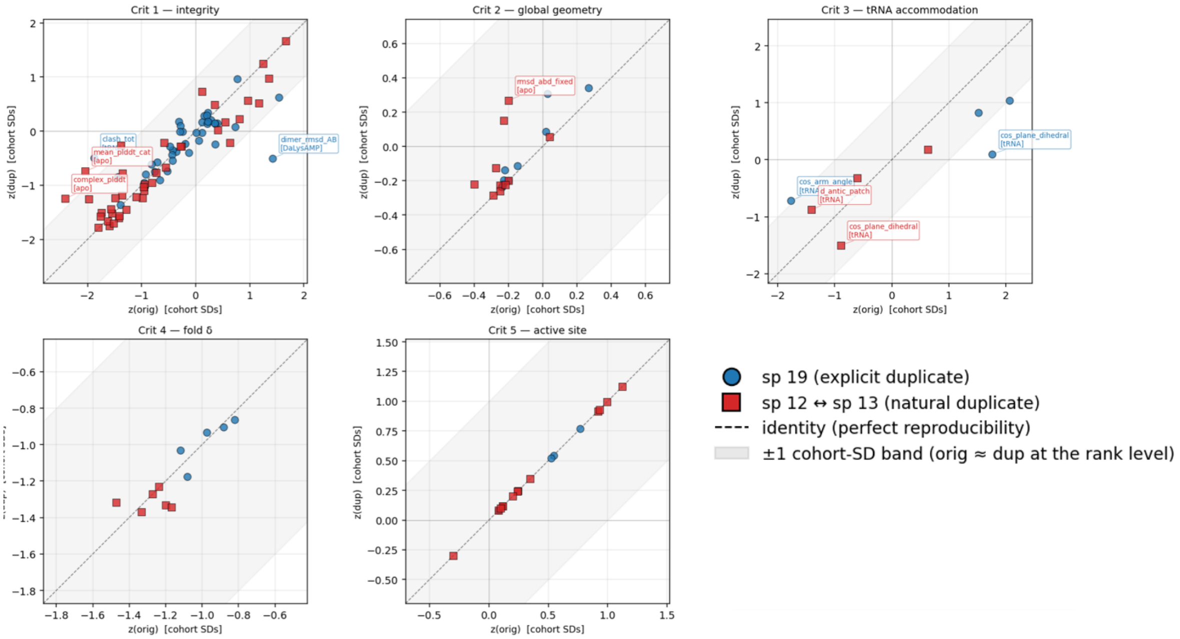


**Figure S4.** Reproducibility analysis of the five criteria. Axes: each ensemble's mean z-scored against the full cohort distribution of per-chimera means for that (metric, config). Diagonal point = the duplicate run would put the chimera in the SAME cohort-relative position (i.e. it would contribute the same z to the composite). Off-diagonal = the chimera's score depends on which Boltz run you happen to read. For criteria 1 and 3, batch effects of >1SD occur, but while criterion 1 measures many parameters, and only few are just beyond acceptable, criterion 3 stats are based on n=10, and were considered more concerning. Consequently, criterion 3's underlying models (tRNA complexes) were run with n=40.

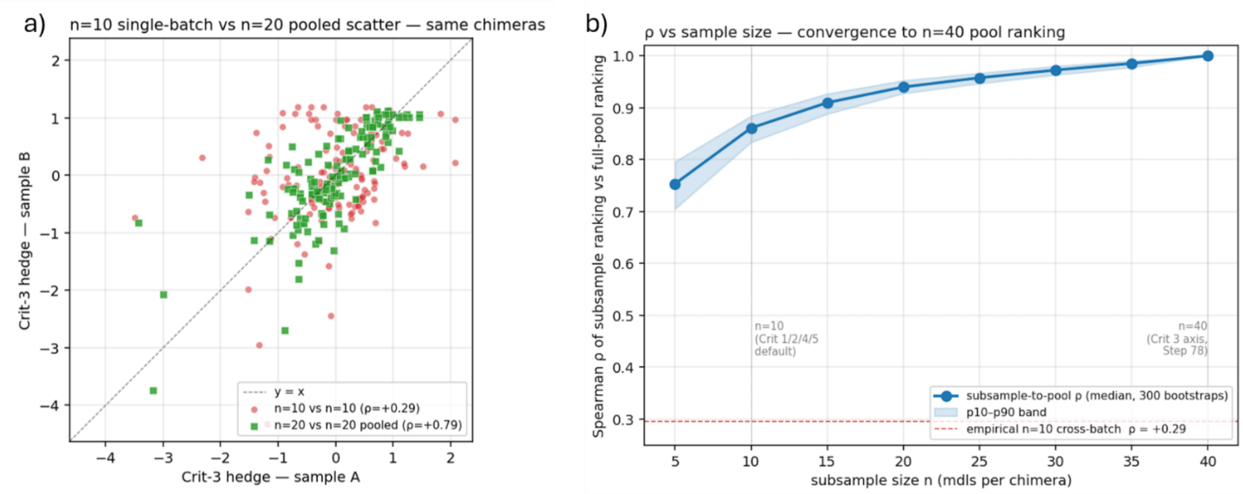


**Figure S5.** Criterion 3 rank reproducibility tightens with n. a) Per-chimera Crit-3 hedge score, sample A vs sample B, for the same set of chimeras at two sampling regimes. Single-batch n = 10 vs n = 10 (red circles, ρ = +0.29) shows substantial scatter off the y = x diagonal; pooling two independent n = 10 batches into n = 20 (green squares, ρ = +0.79) markedly tightens the agreement. Together, the two panels motivate the n = 40 pooled-batch sampling adopted for the two tRNA-containing configurations underlying criterion 3. b) Spearman ρ of the criterion 3 hedge ranking computed on a random subsample of n models per chimera n = 40, as a function of n. Median over 300 bootstrap resamples; shaded band shows the p10–p90 interval. At n = 10 the subsample-to-pool ρ is only ≈ 0.86 and rises to ≈ 0.95 by n = 20 and ≈ 0.97 by n = 30 before saturating at the n = 40 anchor. The empirical cross-batch ρ between two independent n = 10 Boltz batches of the same chimeras (red dashed line, ρ = +0.29) sits well below the within-pool bootstrap band, indicating that batch-to-batch variability dominates over within-batch subsampling noise at n = 10.

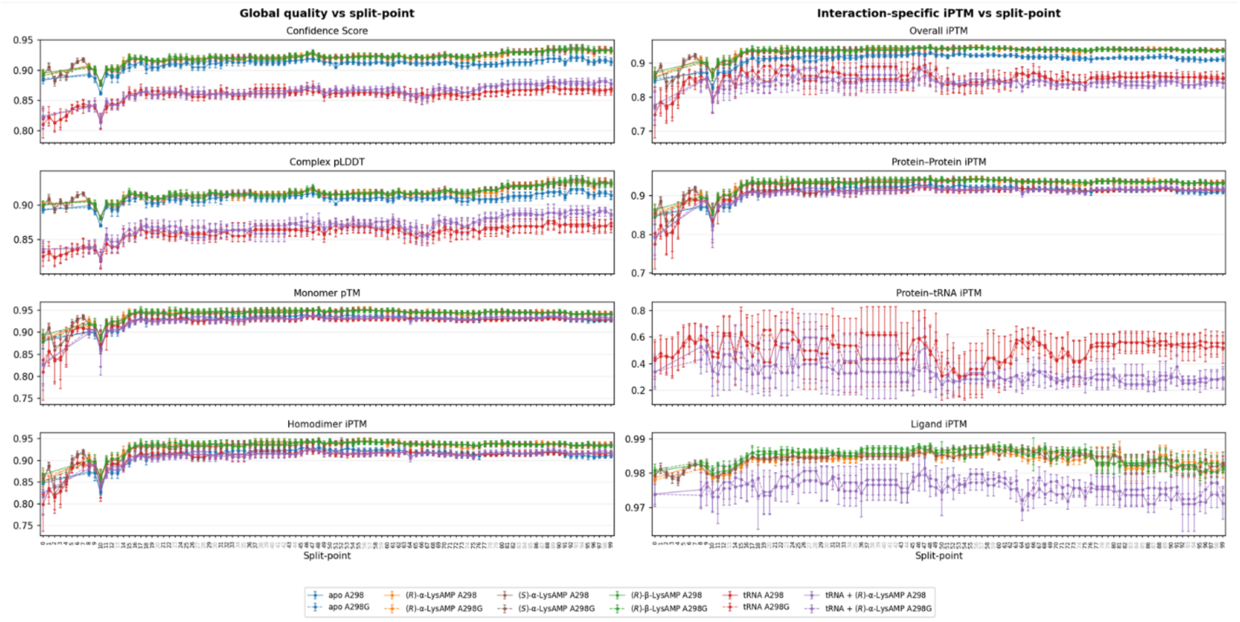


**Figure S6.** Boltz-2 confidence metrics across the chimera split-point sweep. Left column, global model-quality metrics: overall confidence score (Boltz-2 composite), complex-wide mean pLDDT, monomer pTM, and homodimer interface pTM (ipTM). Right column, interaction-specific ipTM: overall ipTM, protein–protein ipTM, protein–tRNA ipTM (tRNA-containing configurations only), and ligand ipTM (ligand-containing configurations only). Lines are per-config means over the model ensemble per (sp, variant) cell; error bars = ±1 SD. Solid lines = A298 variant, dashed = A298G. All global metrics rise sharply over sp0–10 then plateau at high confidence (≈ 0.9), with the tRNA configurations consistently below the protein–protein-dominated configurations. The protein–tRNA ipTM panel shows tRNA-alone (red) and tRNA + R-α-Lys-AMP (purple) tracking in the 0.3–0.6 band, reflecting the lower reliability of cofolding models for protein–nucleic-acid interfaces and motivating the n = 40 sampling for these configurations.

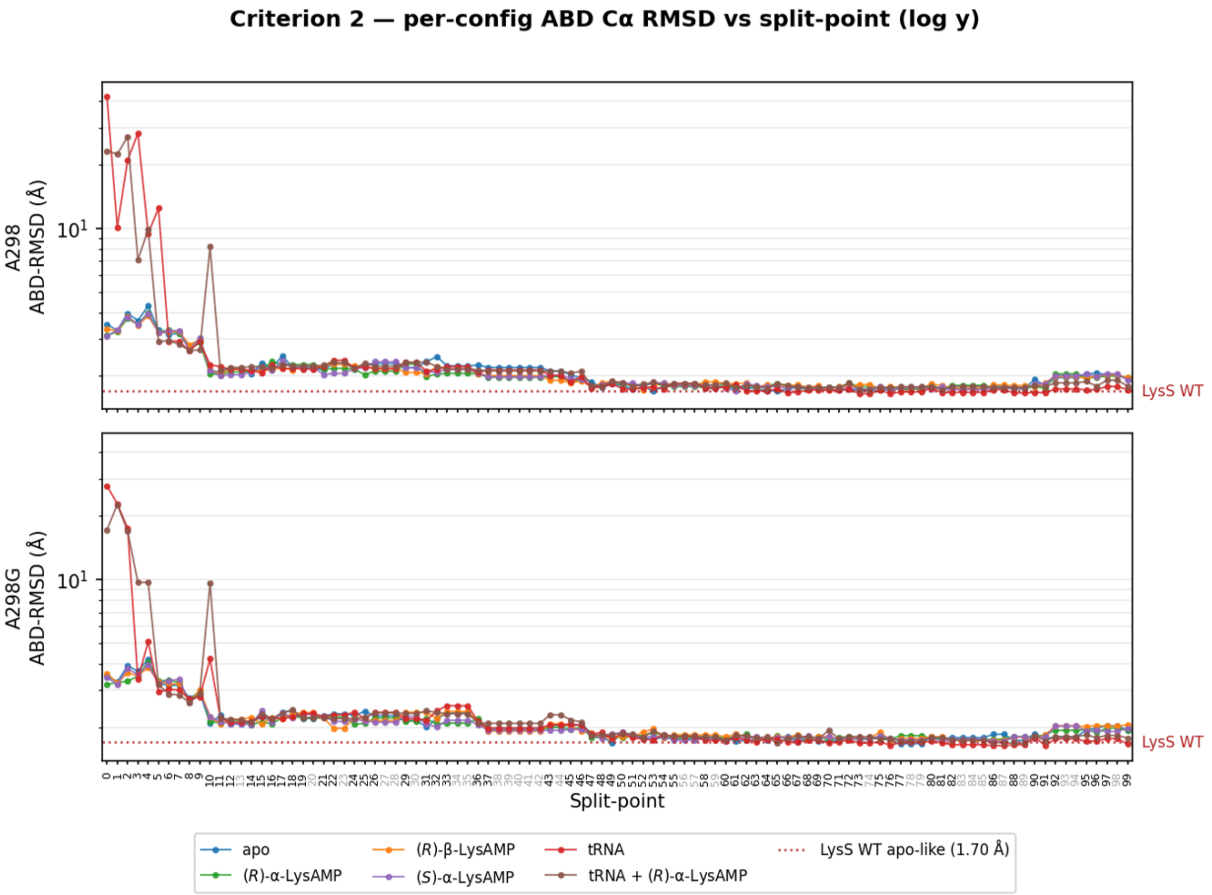


**Figure S7.** Per-configuration Crit-2 ABD Cα-RMSD across the chimera split-point sweep (log y-axis). Top panel: A298 variant; bottom panel: A298G. Each line is the per-(sp, variant, config) mean ABD Cα-RMSD (n = 10 models per config) of the chimera against the 9DPL ABD reference over the fixed 132-pair correspondence map. The criterion-2 composite (fed to the cross-criterion ranking) is the mean across the four apo-like configurations and the gate-passing tRNA-bound models, then cohort-z-scored (sign-flipped; lower = better). Red dotted line: LysS-WT apo-like baseline (1.70 Å). All configurations agree closely from sp≥ 10 onward, sitting just above the WT noise floor. The catastrophic regime at sp0–10 is visible as large excursions in the tRNA and tRNA + R-α-Lys-AMP cells (compressed by the log axis), reflecting failed tRNA docking in the lowest-sp chimeras. Grey-labeled split points are SHEBA sequence-duplicates whose data are inherited from their canonical sp.

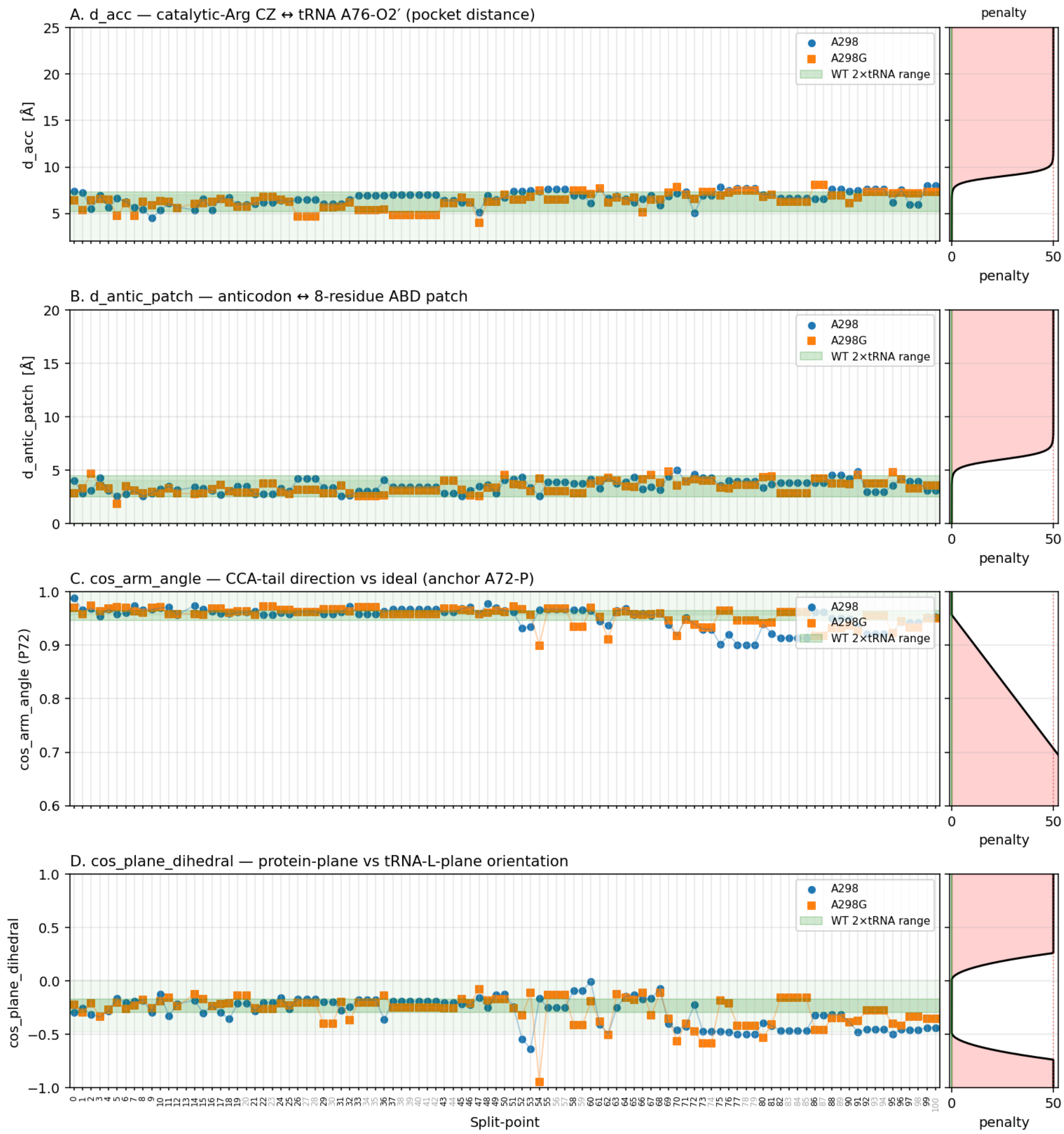


**Figure S8.** Per-submetric Crit-3 tRNA-accommodation values across the chimera split-point sweep and penalty functions. Each panel shows one of the four geometric quantities feeding the per-model productivity score (A: d_acc, catalytic-Arg CZ ↔ tRNA A76-O2′ pocket distance; B: d_antic_patch, anticodon-loop ↔ 8-residue ABD anchor patch; C: cos_arm_angle, CCA-arm direction vs ideal anchor; D: cos_plane_dihedral, protein-plane vs tRNA-L-plane orientation). Points are per-(sp, variant) means over the n = 40 tRNA + R-α-Lys-AMP ensemble; blue = A298, orange = A298G. Green bands mark the wild-type observed range; the lighter-green extension marks the zero-penalty region of the per-model productivity score. Right panels show the penalty function for each metric's axis (red = penalty magnitude): sigmoidal in the distance metrics (A, B), linear in cos_arm_angle (C), and quadratic on either side of the WT window in cos_plane_dihedral (D). Most chimeras across the sp range remain inside the zero-penalty region on all four axes; modest deterioration appears past sp~50 on cos_arm_angle and cos_plane_dihedral.

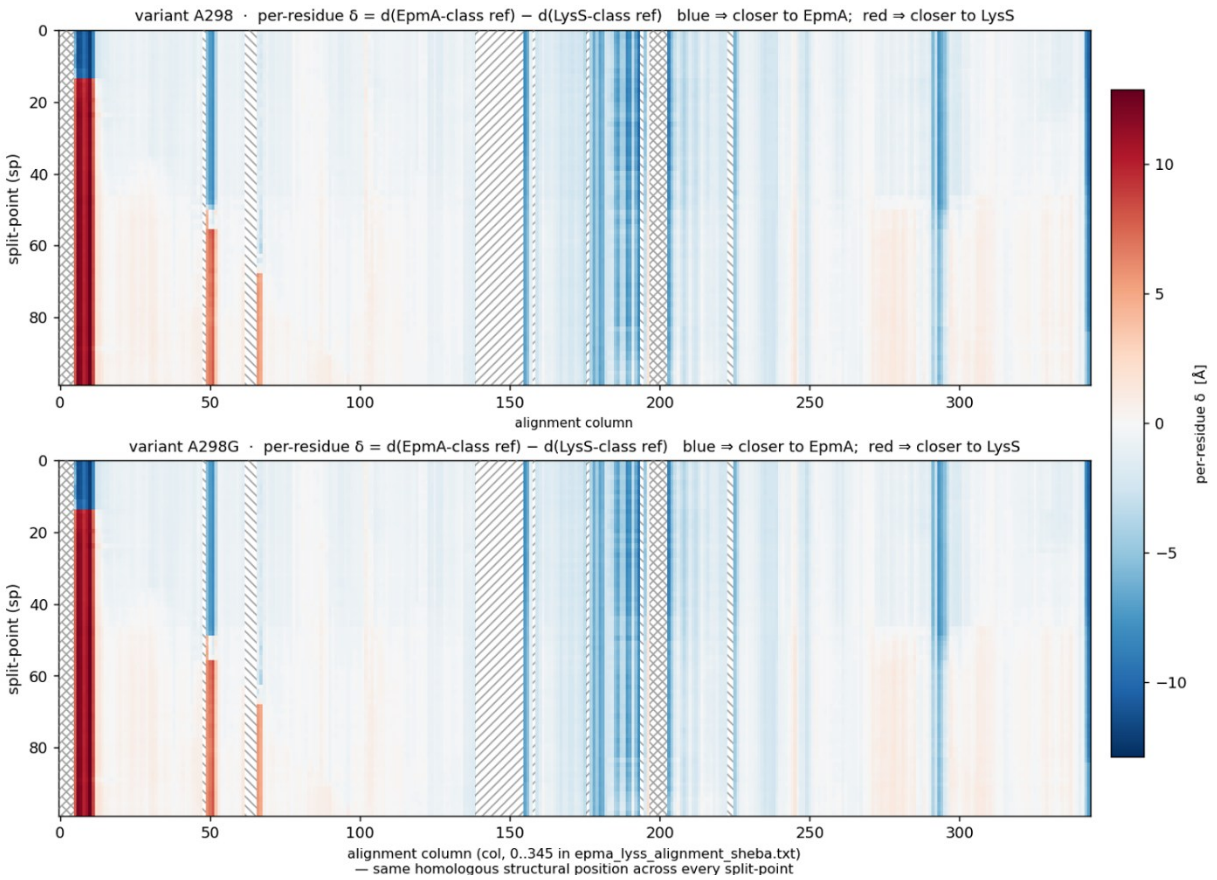


**Figure S9.** Per-residue Crit-4 catalytic-fold drift δ across the (split-point, alignment-column) grid. Heatmaps of the per-residue scalar $\delta = d_{i,\mathrm{LysS}} - d_{i,\mathrm{EpmA}}$, computed from chimera ↔ parent Cα distances pooled across the 6 ligand configurations × n = 10 model ensemble (60 measurements per cell). EpmA-class reference: 3G1Z (apo/tRNA configs) or 3A5Y (holo configs); LysS-class reference: 1BBW (apo/tRNA) or 1E1T (holo). Blue indicates a residue closer to EpmA, red closer to LysS. The residue-summed δ over the catalytic-domain columns is the per-chimera catalytic-fold scalar fed to the composite ranking. Top: native A298 variant. Bottom: A298G specificity-flip mutant. White-hatched columns mark alignment positions where the chimera has a gap or the relevant reference crystal lacks resolved atoms (see legend). Most catalytic-domain columns sit at low |δ| (light shading); localized LysS-drift hotspots (red bands) appear predominantly at the N-terminus, around alignment column ~50, and in the active-site loop region ~190–200; the same residue groups identified as per-residue drift drivers in the manuscript's Crit-4 analysis. The A298 and A298G heatmaps are visually nearly identical, consistent with the broader observation that the A298 ↔ A298G point mutation does not perturb the global catalytic-domain geometry.

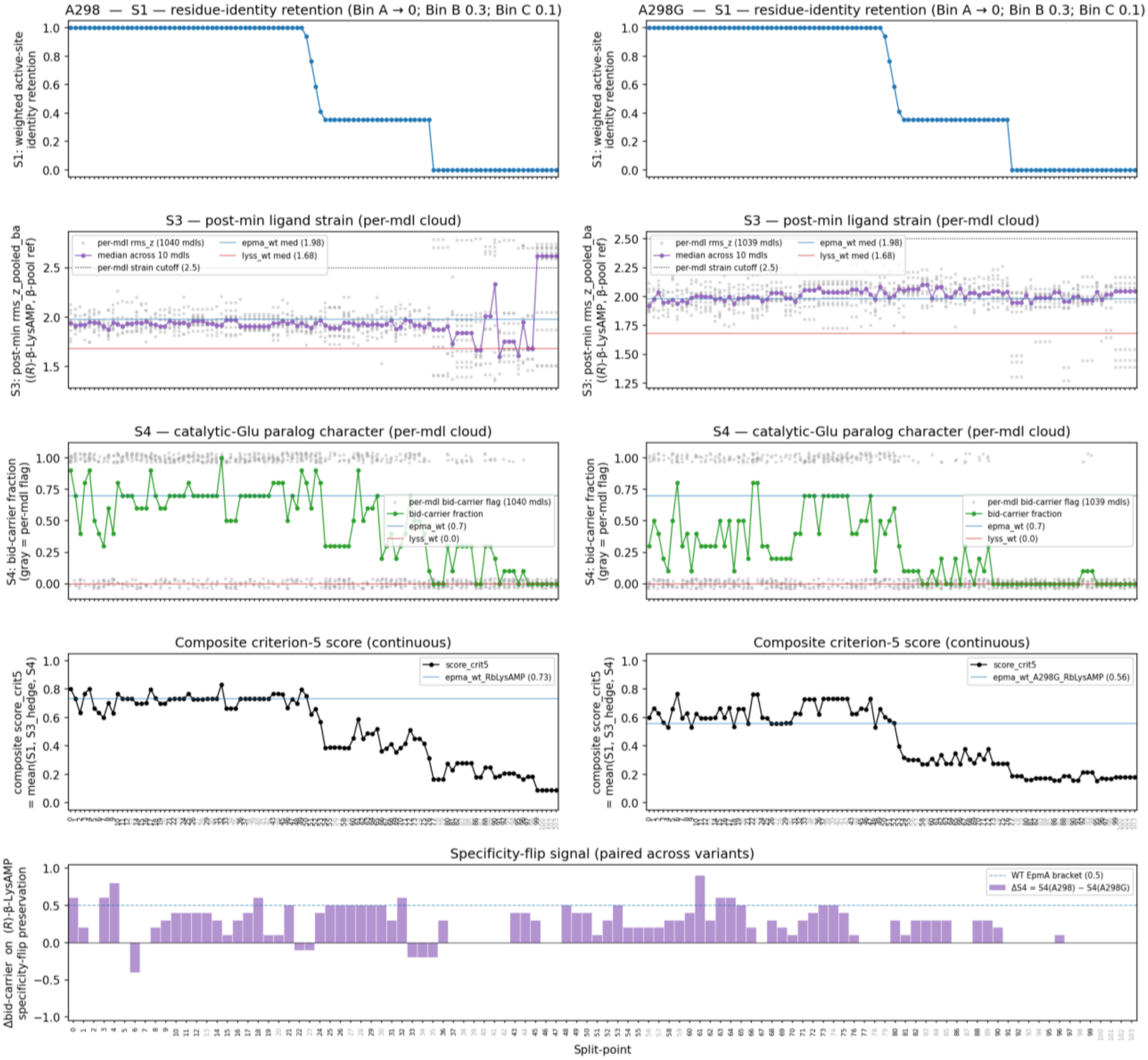


**Figure S10**. Criterion 5 active-site competence per sp, post minimization. Left column: A298; right: A298G. All sub-scores were evaluated on the *(R)*-β-Lys-AMP-bound configuration. Row 1, S1 (AS-identity retention): weighted active-site sequence retention over 11 EpmA residues. Row 2, S3 (post-min ligand strain): per-model rms_z of the AMBER14-minimized (R)-β-Lys-AMP ligand against PDB-pooled β-substrate bond/angle distributions. Grey dots: per-model values (n = 10 per cell); purple line: ensemble median; blue/red lines: EpmA-WT and LysS-WT medians; dotted line: per-model strain cutoff (2.5) used by the hedge pass-rate. Row 3, S4 (catalytic-Glu bid-carrier fraction): per-model carrier-bidentate flag (grey: 0/1 per model); green line: per-cell bid-carrier fraction. The signal stays close to the EpmA-WT reference (0.7) up to sp≈ 70, then collapses toward the LysS-WT reference (0.0) as the catalytic Glu loses its EpmA-type β-clamp. Row 4, composite Crit-5 score: unweighted mean of S1, S3-hedge, and S4. Blue line: WT-EpmA. Bottom panel, specificity-flip signal: Δbid-carrier = bid-carrier(A298) − bid-carrier(A298G), per sp. Dashed line: WT-EpmA. Positive Δ values indicate a chimera preserving the A298-A298G specificity-flip.

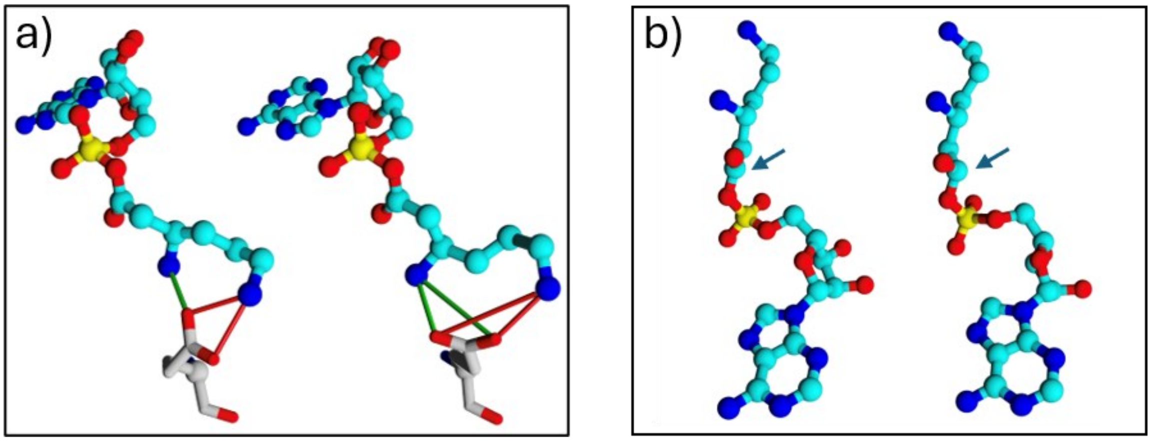


**Figure S11.** Example 3d structures illustrating the criterion 5 geometries. a) The observed bidentate difference of the active site Glu (Glu116 in EpmA, Glu279 in LysS) in the two enzymes for models with *(R)*-β-Lys-AMP ligands. In LysS, the Glu is facing the terminal amine of lysine, while in EpmA models (right), the Glu rotates more toward the beta amine and does not face any amine preferentially anymore. b) Example of a ligand with unrealistic strain. Left: No strain. Right: carbonyl carbon unrealistically (for sp2 hybridization) out of plain.

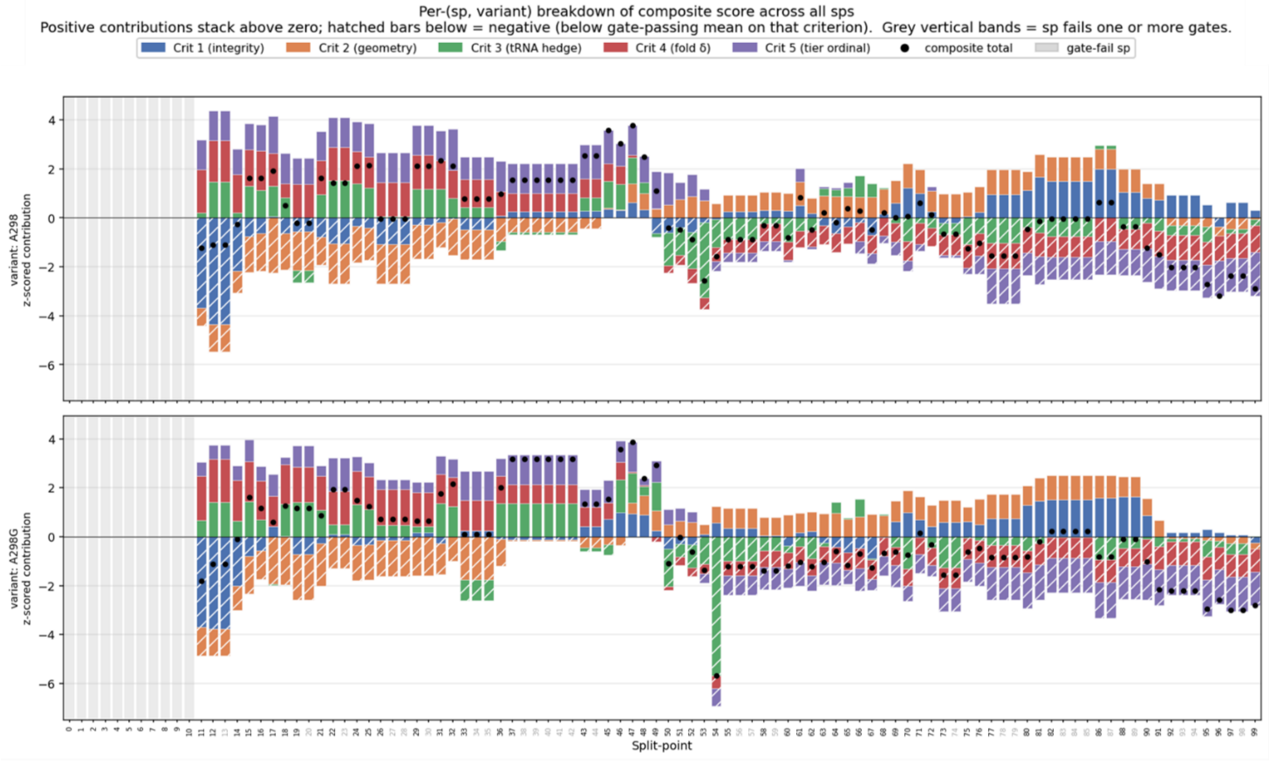


**Figure S12.** Per-(sp, variant) decomposition of the composite score into its five per-criterion z-scored contributions. Top: A298. Bottom: A298G. Stacked bars show each criterion's signed contribution (positive above zero, hatched below = negative); black dots mark the composite total. Grey vertical bands flag chimeras failing one or both gates (Crit 1/Crit 2), excluded from cohort z-scoring.

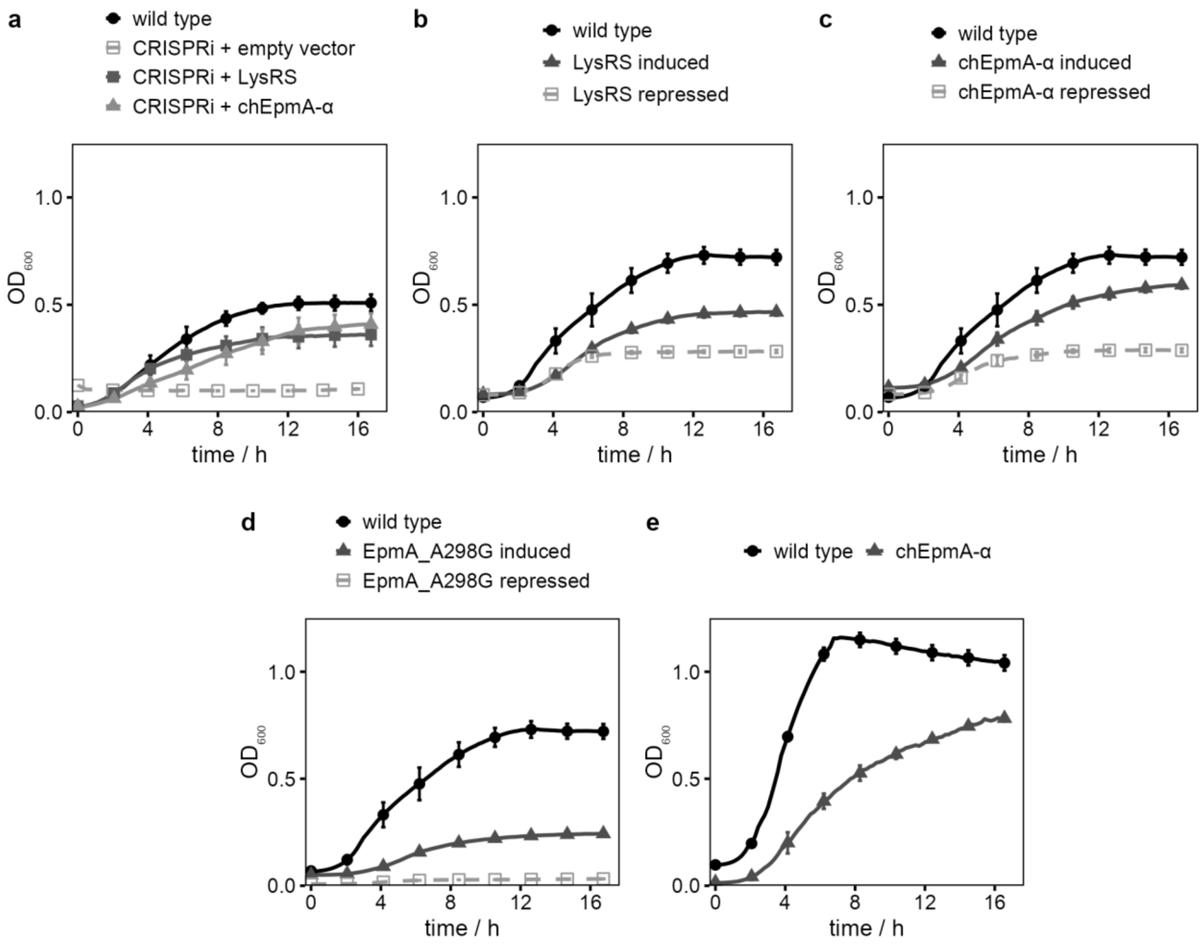


**Figure S13.** Growth complementation assays supporting replacement of LysRS by engineered chEpmA variants. *E. coli* strains were grown in LB at 37 °C in a Spark 20M microplate reader (Tecan), and $OD_{600}$ was recorded every 20 min. **(a)** CRISPRi knockdown of genomic *lysS* in the *E. coli* Δ*lysU* background, comparing wild type, CRISPRi with empty vector (no rescue), CRISPRi complemented *in trans* with plasmid-borne LysRS, and CRISPRi complemented with plasmid-borne chEpmA-α. **(b–d)** Plasmid-based complementation in the *E. coli* Δ*lysS* Δ*lysU* background with LysRS **(b)**, chEpmA-α **(c)**, or EpmA_A298G **(d)**, grown in the presence of 0.2% (w/v) L-(+)-arabinose (induced) or 0.2% (w/v) glucose (repressed); residual growth under repressed conditions reflects basal (leaky) expression from the P_BAD promoter. **(e)** Growth of the *E. coli* Δ*lysS* Δ*lysU* strain relying exclusively on a genomically integrated copy of chEpmA-α, compared with wild type, grown without antibiotic/inducer selection. Data are shown as mean ± SD of 3 biological replicates. Statistical significance was assessed by unpaired two-tailed Student's *t*-test (ns, not significant; *p < 0.05; **p < 0.01; ***p < 0.001). 'wild type' denotes the unmodified parental strain (a, e) or the parental strain carrying the respective empty vector (b–d).

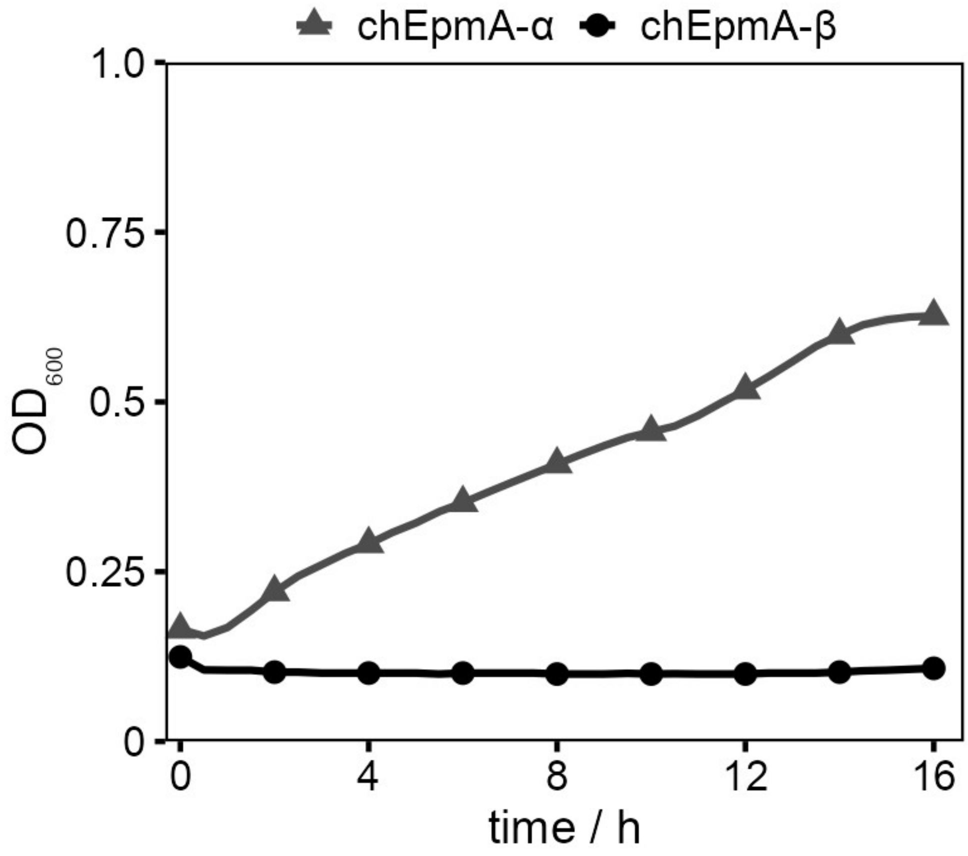


**Figure S14.** *In vivo* toxicity associated with expression of the β-lysine-specific chimera chEpmA-β. *E. coli* Δ*lysS* Δ*lysU* cells carrying plasmid-borne chEpmA-α (A298G, (*S*)-α-lysine-specific) or chEpmA-β (A298, (*R*)-β-lysine-specific) were grown in LB at 37 °C in the presence of 0.2% (w/v) L-(+)-arabinose to induce expression; $OD_{600}$ was recorded every 20 min in a Spark 20M microplate reader (Tecan). Curves represent the mean of 3 independent biological replicates. While chEpmA-α sustains growth, expression of chEpmA-β abolishes growth, consistent with proteome-wide misincorporation of (*R*)-β-lysine at canonical lysine codons following mischarging of the endogenous $tRNA^{Lys}$ pool.

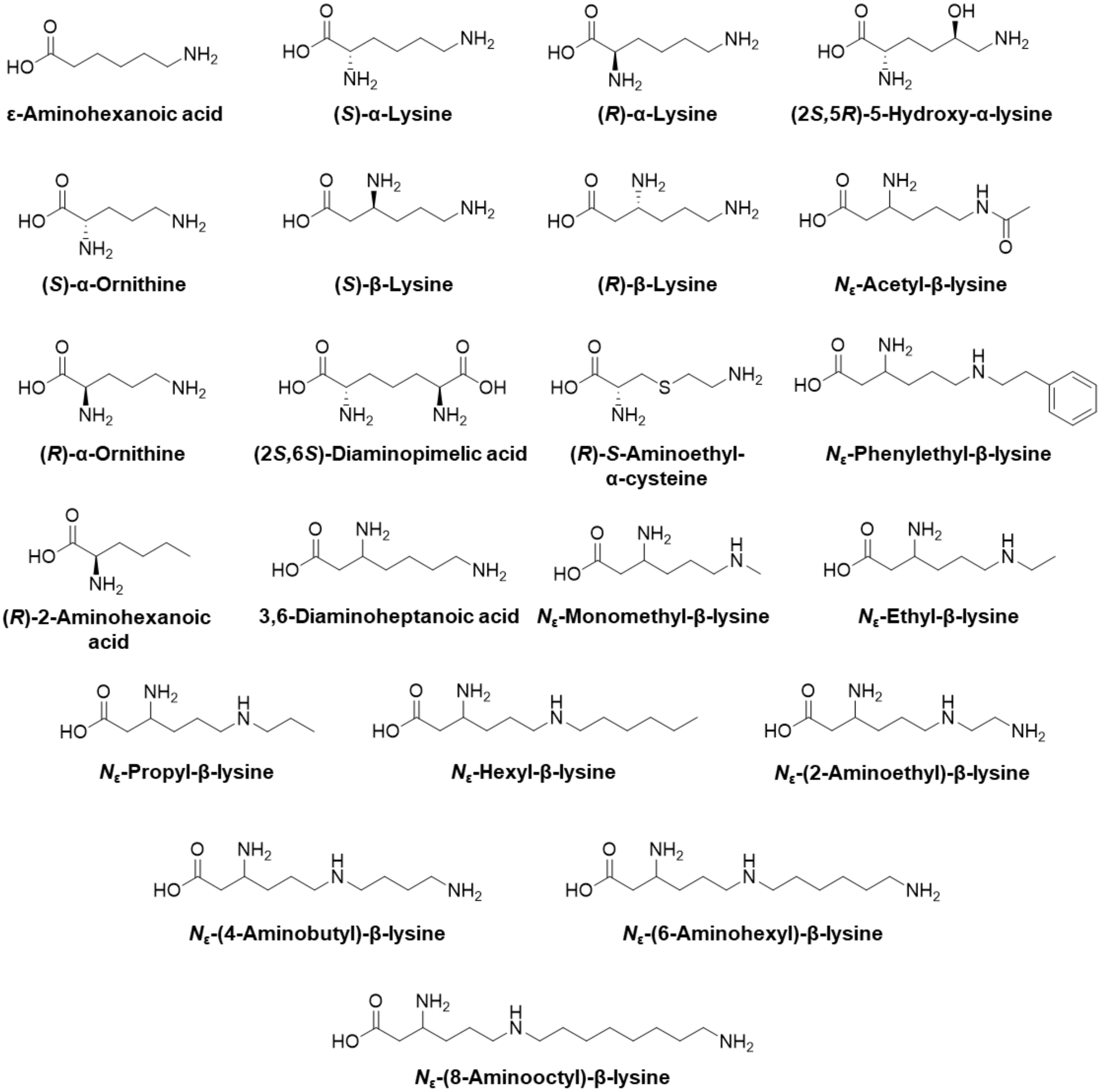


**Figure S15.** The reported donor substrate spectrum of EpmA and its variant EpmA_A298G Stereoselectivity is given, where possible [2].

#### Supplementary Tables

#### Table S1. Bacterial strains used in this study.

| Strain | Relevant genotype | Purpose | Resistance | Reference |
|---|---|---|---|---|
| *E. coli* DH5α λ*pir* | *endA1 hsdR17 glnV44 thi-1 recA1 gyrA96 relA1* φ80d*lac*ΔM15 Δ(*lacZYA-argF*)U169 *zdg-232*::Tn*10 uidA*::*pir*$^{+}$ | Plasmid cloning | – | [13] |
| *E. coli* LMG194 | F$^{-}$ Δ(*lacIPOZY*)X74 *galE galK thi rpsL* Δ*phoA ara714* | Protein production | – | [14] |
| *E. coli* BW25113 | F$^{-}$ λ$^{-}$ Δ(*araD-araB*)567 Δ*lacZ*4787(::*rrnB*-3) *rph-1* Δ(*rhaD-rhaB*)568 *hsdR514* | Parent strain | – | [15] |
| BW25113 Δ*lysS* (JW2858-4) | BW25113; Δ*lysS*741::*kan* | Strain construction | $Km^{R}$ | [15] |
| BW25113 Δ*lysU* (JW4090-1) | BW25113; Δ*lysU*756::*kan* | Strain construction | $Km^{R}$ | [15] |
| BW25113 Δ*lysS* Δ*lysU* | BW25113; Δ*lysS* Δ*lysU* | Complementation assays | – | This study |
| BW25113 Δ*lysS* Δ*lysU* Δ*epmA*::*chEpmA-α* | BW25113 Δ*lysS* Δ*lysU*; genomic *epmA* replaced by *chEpmA*-α (sp43, A298G) | Genome-integration validation | – | This study |
| LC-E75 | *E. coli* MG1655; pOSIP-CO-RBS-library-*dCas9* at 186 *attB*; pOSIP-KL-mCherry at λ *attB* | CRISPRi parent strain | – | [16] |
| LC-E75 Δ*lysU* | LC-E75; Δ*lysU* | CRISPRi knockdown of *lysS* | – | This study |

**Table S2. Oligonucleotides used in this study.**

| Primer name | Sequence (5′→3′) |
|---|---|
| sgRNA_lysS_1 | GCT CAG TCC TAG GTA TAA TAC TAG TCA AAG CAC TGC GTC CGC TGC GTT TTA GAG CTA GAA ATA GCA |
| sgRNA_lysS_2 | CAC TTT TTC AAG TTG ATA ACG GAC TAG CCT TAT TTT AAC TTG CTA TTT CTA GCT CTA AAA CGC AG |
| sgRNA_lysS_3 | AGT CCG TTA TCA ACT TGA AAA AGT GGC ACC GAG TCG GTG CTT TTT TTG AAT TCT CTA GAG TCG AC |
| lysS_CRISPRi_FW | TAG TCA TTA AAG ATC TGC GCA CCG GGG |
| lysS_CRISPRi_RV | AAA CCC CCG GTG CGC AGA TCT TTA ATG |
| Amp_sgRNA_FW | GCT CAG TCC TAG GTA TAA TAC TAG T |
| Amp_sgRNA_RV | GTC GAC TCT AGA GAA TTC AAA AAA AA |
| pTarget_BB1_FW | TCT CAT GAC CAA AAT CCC TTA ACG TGA GTT TTC GTT CCA CTG AGC |
| pTarget_BB1_RV | ACT AGT ATT ATA CCT AGG ACT GAG CTA GCT GTC AAG GAT CCA GCA |
| pTarget_BB2_FW | TTT TTT TGA ATT CTC TAG AGT CGA CCT GCA GAA GCT TAG ATC TAT |
| pTarget_BB2_RV | GCT CAG TGG AAC GAA AAC TCA CGT TAA GGG ATT TTG GTC ATG AGA |
| UHA_lysS_FW | GTA AAA GCC CGT TTG ACT CC |
| UHA_lysS_RV | GCG CCC TGT GCG TGT TGT TCA GAC ATG TTG GTT CCT CAT AAC CCT |
| Hybrid_FW | AGG GTT ATG AGG AAC CAA CAT GTC TGA ACA ACA CGC ACA GGG CGC |
| Hybrid_RV | GGT TGT GAG CAT AAC GTA ATG CTT ATG CCC GGT CAA CGC TAA AGG |
| DHA_FW | CCT TTA GCG TTG ACC GGG CAT AAG CAT TAC GTT ATG CTC ACA ACC |
| DHA_RV | TCT CGC GTT CTC CAG TTG GC |
| SacI_sRBS_lysS_FW | CGC GAG CTC GAA AAT AAA TAA ATT GCT AGT AAC GGC AGT CTT TAA TAA GGA GGT TTT TTA TGT CTG AAC AAC ACG CAC AGG |
| lysS186_EpmA_OL_RE | CGC CGC GCG CAC TTT AAA GGT GTT GCG GGA T |
| lysS186_EpmA_OL_FW | CTT TAA AGT GCG CGC GGC GAT TAT GG |
| lysS210_EpmA_OL_RE | CTG GCT CAT CAT CGG CGT TTC AAC TTC CAT AA |
| lysS210_EpmA_OL_FW | GAA ACG CCG ATG ATG AGC CAG GCG ACG GTA A |
| SacI_lysS_EpmA_Hyb_FW | GGA TCC GAG CTC TCT GAA CAA CAC GCA CAG GGC |
| KpnI_lysS_EpmA_Hyb_RV | GCT GGT ACC TTA TGC CCG GTC AAC GCT AAA G |
| XbaI_lysS_GS-His6_RV | CGA CTC TAG ATT AGT GAT GGT GAT GGT GAT GGC TGC CTT TTT ACC GGA CGC ATC GCC G |
| A298G_FW | CAG CAG GTC TGG CGA TTG GTA TCG |
| A298G_RV | CGA TAC CAA TCG CCA GAC CTG CTG |
| lysS_chk_FW | GCC AGA TTC GTT CTT ATG TCC T |
| lysS_chk_RV | TCA GTC CGT GGA TGG TGA TG |
| lysU_chk_FW | CGA GGC AGA TAA TAC AGT GTA C |
| lysU_chk_RV | TAG GAG CTG TAG TTG AGT GGA C |
| kan_FW | GGC AAT CAG GTG CGA CAA TC |
| kan_RV | CGA TTC CGA CTC GTC CAA CA |

#### Table S3. Plasmids used in this study.

| Plasmid | Description | Backbone | Resistance | Induction | Reference |
|---|---|---|---|---|---|
| psgRNAc | Expresses sgRNA; p15A low-copy origin | pUC19 | $Cm^R$ | – | [16] |
| pRED/ET® | λ-RED recombinase for gene deletion | pBAD24 | $Cb^R$ | Arabinose | Gene Bridges GmbH, Heidelberg, Germany |
| 709-FLPe | FLPe recombinase for FRT-cassette removal | – | $Cb^R$ | 30 °C | Gene Bridges GmbH, Heidelberg, Germany |
| pBAD33_*lysS* | sRBS::*lysS*–$His_6$ (C-terminal) | pBAD33 | $Cm^R$ | Arabinose | This study |
| pBAD33_chEpmA-α (sp43) | sRBS::*lysS*1–186–*epmA*_A298G–$His_6$ (sp43) | pBAD33 | $Cm^R$ | Arabinose | This study |
| pBAD33_chEpmA-β (sp43) | sRBS::*lysS*1–186–*epmA*_A298–$His_6$ (sp43) | pBAD33 | $Cm^R$ | Arabinose | This study |
| pBAD33_*epmA*_A298G | sRBS::$His_6$–*epmA*_A298G | pBAD33 | $Cm^R$ | Arabinose | This study |
| pCas9 | Constitutive Cas9 expression and arabinose-inducible λ-RED | pKD46 | $Kan^R$ | Arabinose; | [1] |
| pTargetF | Constitutive sgRNA expression without target | Trc99a | SpecR | – | [3] |
| pTarget_*lysS* | pTargetF derivative expressing sgRNA targeting *lysS* | Trc99a | SpecR | – | This study |

#### Table S4. DNA templates and oligonucleotides for *in vitro* transcription of $tRNA^{Lys}$.

| Template / Primer | Sequence (5′→3′) |
|---|---|
| T7 promoter (primer) | GAA ATT AAT ACG ACT CAC TAT A |
| $tRNA^{Lys}$ template | CTT TAA TTA TGC TGA GTG ATA TCC CAG CAA TCG AGT CAA CCA TCT CGT CAA CTG AAA ATT AGT TAA CCA GCG TCC AAG CTT AGG ACG TGC TGG GTG GT |
| tRNA_OL1 | CCA ATG CAT ACC AAA CCT GCA CGC TAG TTT CCT GAT GAA CA |
| tRNA_OL2 | AAT AGT TTA ATT ATC GAC AGA GGT GTA ATT GCT GGA AAA TGT TCA TCA GGA AAC TAG CGT |
| tRNA_OL3 | ACA CCC TGT CGA TAA TTA ACT ATT TGA CGA AAA GCT GAA AAC CAC TAG AAT GCG CCT CCG |

| Template / Primer | Sequence (5′→3′) |
|---|---|
| tRNA_OL4 | CCA CCG CTT CAT ACC ATC AAT TCT TAA AAA GAA TTG GTA CCA CGG AGG CGC ATT CTA GTG |
| tRNA_OL5 | ATT GAT GGT ATG AAG CGG TGG CTC AAT GGT AGA GCT TTC GAC TCT AAA TCG AAG GGT TTG C |
| tRNA_OL6 | CGG GGT TTC AAA ATT GGT GAA ACG GAC AGG AAT TGA ACC TGC AAC CCT TCG ATT TAG AGT C |
| tRNA_OL7 | TTT CAC CAA TTT TGA ACC CCG CTT CGG CGG GGT TTT TTG TTT TTC TGT GCA TTT CGT CAC C |
| tRNA_OL8 | CCA TAT TAC GGG CGT TTA TTT GCG AAG GGG AGG GTG ACG AAA TGC ACA GAA AAA |

#### References


[1] M. Hellwig, S. Geissler, R. Matthes, A. Peto, C. Silow, M. Brandsch, T. Henle, “Transport of Free and Peptide-Bound Glycated Amino Acids: Synthesis, Transepithelial Flux at Caco-2 Cell Monolayers, and Interaction with Apical Membrane Transport Proteins” *ChemBioChem* **2011**, *12*, 1270–1279.

[2] L. Cui, A. Vigouroux, F. Rousset, H. Varet, V. Khanna, D. Bikard, “A CRISPRi screen in E. coli reveals sequence-specific toxicity of dCas9” *Nat. Commun.* **2018**, *9*, DOI 10.1038/s41467-018-04209-5.

[3] Y. Jiang, B. Chen, C. Duan, B. Sun, J. Yang, S. Yang, “Multigene editing in the Escherichia coli genome via the CRISPR-Cas9 system” *Appl. Environ. Microbiol.* **2015**, *81*, 2506–2514.

[4] M. van Kempen, S. S. Kim, C. Tumescheit, M. Mirdita, J. Lee, C. L. M. Gilchrist, J. Söding, M. Steinegger, “Fast and accurate protein structure search with Foldseek” *Nat. Biotechnol.* **2024**, *42*, 243–246.

[5] J. Jung, B. Lee, *Protein structure alignment using environmental profiles*, **2000**.

[6] S. Henikoff, J. G. Henikoff, *Amino acid substitution matrices from protein blocks (amino add sequence/alignment algorithms/data base srching)*, **1992**.

[7] P. J. A. Cock, T. Antao, J. T. Chang, B. A. Chapman, C. J. Cox, A. Dalke, I. Friedberg, T. Hamelryck, F. Kauff, B. Wilczynski, M. J. L. De Hoon, “Biopython: Freely available Python tools for computational molecular biology and bioinformatics” *Bioinformatics* **2009**, *25*, 1422–1423.

[8] G. Landrum, P. Tosco, B. Kelley, R. Rodriguez, D. Cosgrove, R. Vianello, sriniker, P. Gedeck, G. Jones, D. Nealschneider, E. Kawashima, N. Schneider, tadhurst-cdd, A. Dalke, M. Swain, B. Cole, S. Turk, A. Savelev, N. Maeder, Y. Pechersky, A. Vaucher, M. Wójcikowski, R. Walker, H. Faara, I. Take, V. F. Scalfani, D. Probst, K. Ujihara, J. Monat, J. Lehtivarjo, "RDKit: Open-source cheminformatics," **2028**.

[9] J. Wohlwend, G. Corso, S. Passaro, M. Reveiz, K. Leidal, W. Swiderski, T. Portnoi, I. Chinn, J. Silterra, T. Jaakkola, R. Barzilay, "Boltz-1 Democratizing Biomolecular Interaction Modeling." *bioRxiv* **2024**, DOI 10.1101/2024.11.19.624167.

[10] M. Mirdita, K. Schütze, Y. Moriwaki, L. Heo, S. Ovchinnikov, M. Steinegger, "ColabFold: making protein folding accessible to all" *Nat. Methods* **2022**, *19*, 679–682.

[11] J. A. Maier, C. Martinez, K. Kasavajhala, L. Wickstrom, K. E. Hauser, C. Simmerling, "ff14SB: Improving the Accuracy of Protein Side Chain and Backbone Parameters from ff99SB" *J. Chem. Theory Comput.* **2015**, *11*, 3696–3713.

[12] E. Krieger, K. Joo, J. Lee, J. Lee, S. Raman, J. Thompson, M. Tyka, D. Baker, K. Karplus, **2009**, DOI: 10.1002/prot.22570.

[13] E. Krieger, G. Koraimann, G. Vriend, "Increasing the precision of comparative models with YASARA NOVA - A self-parameterizing force field" *Proteins: Structure, Function and Genetics* **2002**, *47*, 393–402.

[14] E. Krieger, G. Vriend, "YASARA View - molecular graphics for all devices - from smartphones to workstations" *Bioinformatics* **2014**, *30*, 2981–2982.

[15] T. Baba, T. Ara, M. Hasegawa, Y. Takai, Y. Okumura, M. Baba, K. A. Datsenko, M. Tomita, B. L. Wanner, H. Mori, "Construction of Escherichia coli K-12 in-frame, single-gene knockout mutants: the Keio collection" *Mol. Syst. Biol.* **2006**, *2*, MSB4100050.

[16] L. Cui, A. Vigouroux, F. Rousset, H. Varet, V. Khanna, D. Bikard, "A CRISPRi screen in E. coli reveals sequence-specific toxicity of dCas9" *Nat. Commun.* **2018**, *9*, DOI 10.1038/s41467-018-04209-5.
